# Stack Mapping Anchor Points (SMAP): a versatile suite of tools for read-backed haplotyping

**DOI:** 10.1101/2022.03.10.483555

**Authors:** Dries Schaumont, Elisabeth Veeckman, Felix Van der Jeugt, Annelies Haegeman, Sabine van Glabeke, Yves Bawin, Joanna Lukasiewicz, Sebastien Blugeon, Philippe Barre, Maria de la O Leyva-Pérez, Stephen Byrne, Peter Dawyndt, Tom Ruttink

## Abstract

Here we present SMAP, a software package that implements a suite of computational tools to extract multi-allelic haplotypes using read-backed haplotyping. SMAP tools first perform accurate read processing and analyze read mapping distributions across sample sets. Then, two complementary modules can be invoked for haplotype calling: *SMAP haplotype-sites* combines known Single Nucleotide Polymorphisms (SNPs) and/or read mapping position polymorphisms (SMAPs) to reconstruct compressed, read-reference-encoded haplotype strings. In contrast, *SMAP haplotype-window* works independent of prior knowledge of polymorphisms, groups reads by locus, defines a window enclosed between two custom border sequences, and retains the entire corresponding DNA sequence as haplotype. *Haplotype-window* is, among many applications, especially useful for high-throughput CRISPR/Cas mutation screens. Either way, SMAP creates a single integrated haplotype call table across all loci and samples. SMAP haplotyping is extremely versatile and can be applied to highly multiplex amplicon sequencing (HiPlex), Shotgun (*e.g.* whole genome shotgun (WGS) sequencing, probe capture and RNA-Seq), or Genotyping-by-Sequencing (GBS) data; and to Illumina short reads, PacBio and MinION long reads. SMAP creates discrete genotype calls for individuals of any ploidy or quantitative haplotype frequency spectra for Pool-Seq data, and can scale from tens to thousands of loci and/or samples. SMAP, including the source code written in Python is available at https://gitlab.com/truttink/smap, and a detailed user manual and guidelines for accurate read processing is available at https://ngs-smap.readthedocs.io/, under the GNU Affero General Public License v3.0.

---

Genetic polymorphism is fundamental for genetic analysis and a plethora of approaches based on Next Generation Sequencing (NGS) data have been developed to characterize genetic variation. Until now, many computational tools have focused on discovery of, and genotyping at, SNPs. However, in species with high levels of genetic diversity multiple SNPs can occur within a single sequencing read. Read-backed haplotyping is the process of using NGS read information to arrange genotype calls for neighboring polymorphisms per physical DNA molecule, *i.e.* corresponding to a haploid chromatin string in the nucleus. Phasing neighboring sequence variants into haplotypes creates multi-allelic molecular markers, increasing the molecular marker diversity information content per locus, and thus the discriminative power and the resolution of genetic diversity and population studies (Lu *et al*., 2011; Sallam *et al*., 2020). On the one hand, SNPs in phase carry redundant information, and should be reduced to avoid multiple testing during association genetics or to avoid creating genetic maps with excessive numbers of co-located markers. On the other hand, bi-allelic SNPs may be part of different multi-allelic haplotypes, and confound accurate estimation of the respective allele frequencies in population genetics (**Supplementary Fig. S1**). A major advantage of using short haplotypes over single bi-allelic SNPs is their increased information content, leading to a stronger linkage disequilibrium (LD) with quantitative trait loci (QTL) flanking the marker on the same DNA strand and/or more accurate genomic predictions (*e.g.* Calus *et al*., 2009; Matias *et al*., 2017).

Here we present the SMAP package, a versatile suite of tools for read-backed haplotyping (**Supplementary Fig. S2**). SMAP addresses some of the inherent limitations of existing haplotyping tools that often have a specific scope of application (*e.g.* diploid individuals, viral quasi-species, fixed haplotype block lengths, limited maximal SNP number, short read length, requirement of prior knowledge of variants or genetic relationships; **Supplementary Table S1** and references therein). Improvements covered by SMAP include the optimization of locus delineation based on customized start and end sites per NGS library method (**Fig. 1**, **Fig. 2**, **Fig. 3**), expanding the types and the number of variants per locus, becoming independent from prior knowledge of variants and/or haplotypes (*de novo* discovery of polymorphisms and/or phases), flexible application across all main NGS library types, including short and long read sequencing technologies, and sample types including polyploids and Pool-Seq samples, and scalability from tens to thousands of loci and/or samples (**Fig. 1** and **Supplementary Table S1**).

**Figure 1.**
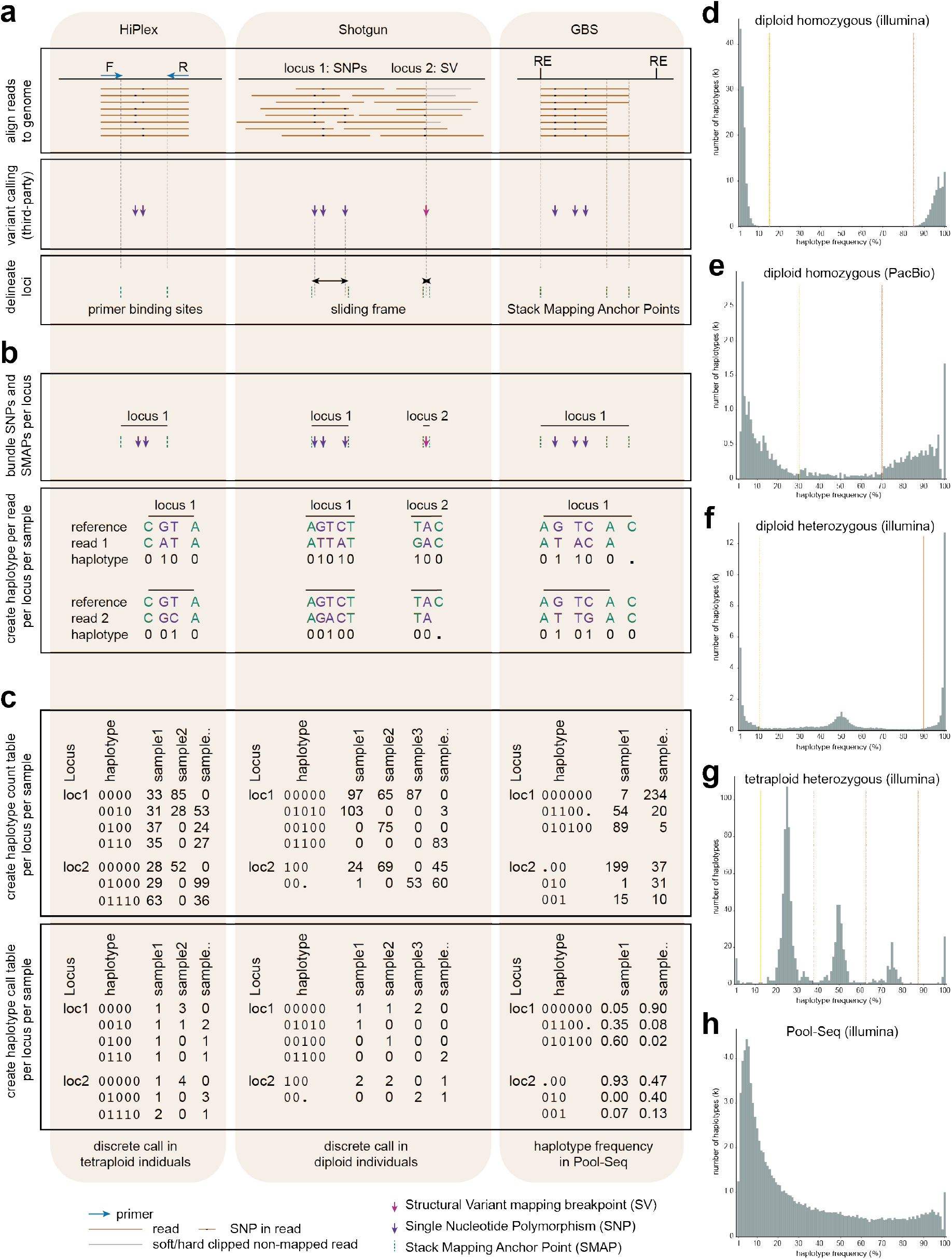
Scheme detailing locus delineation and haplotype calling with *SMAP haplotype-sites*. **a)** locus delineation is based on: primer binding sites (HiPlex); dynamic sliding frames spanning known neighboring SNPs or structural variants (Shotgun); or Stack Mapping Anchor Points (GBS, see details in **Fig. 3** and **Supplementary Fig. S3**). **b)** Read-backed haplotyping using read-reference aligned nucleotide positions. Haplotype strings are encoded as: absent “.”, reference “0”, alternative “1”, or gap “-“ (gaps not shown). **c)** Creation of haplotype count tables and quantification of haplotype frequencies for Pool-Seq (example shown for GBS data) or transformation to discrete dosage calls (using haplotype frequency intervals) in diploid (*e.g.* Shotgun), or tetraploid (*e.g.* HiPlex) data. Haplotype frequency spectra of: **d)** SV deletion calling in WGS Shotgun reads in a homozygous diploid *Oryza sativa* ssp. *japonica* individual; **e)** haplotype calling of SNPs in sliding frames of 1000 bp using PacBio long reads in a homozygous diploid *A. thaliana* Ler individual, (Jiao *et al.*, 2020); **f)** SNP/SMAP haplotyping in *Pst*I-GBS of a heterozygous diploid *L. perenne* individual; **g)** SNP haplotyping in the PotatoMASH HiPlex panel in a tetraploid *S. tuberosum* individual; **h)** SNP/SMAP haplotyping in *Pst*I Pool-Seq GBS of a pool of 48 diploid *L. perenne* individuals.

**Figure 2.**
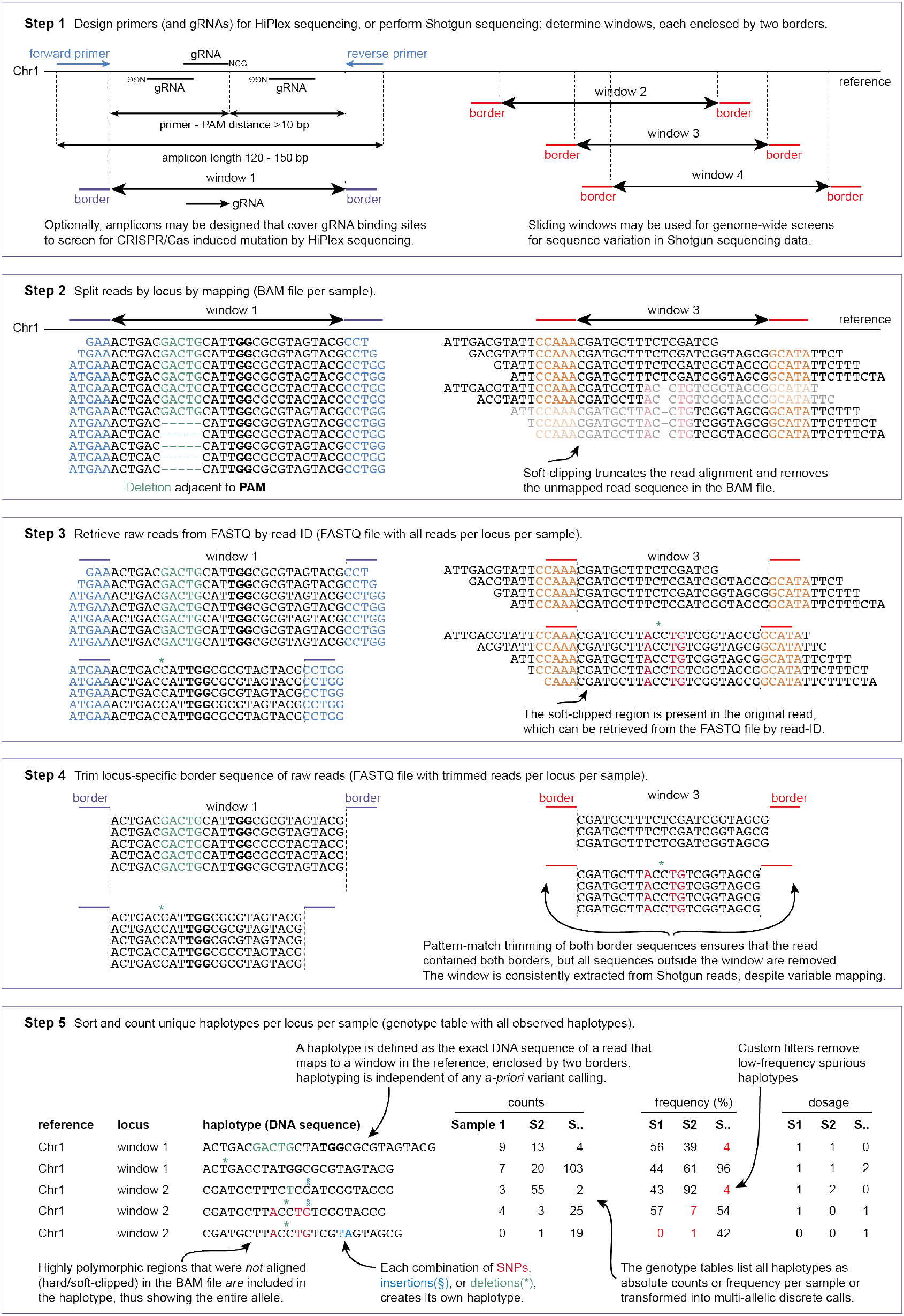
Scheme detailing locus delineation and haplotype calling with *SMAP haplotype-window*. Borders enclosing windows may be defined based on HiPlex amplicon sequencing primers (covering gRNA binding sites for CRISPR/Cas induced mutations), or as sliding windows to analyze Shotgun sequencing data, with fixed or dynamic border length, window length and window distance. *SMAP haplotype-window* can screen for CRISPR/Cas-induced or naturally occurring mutations, independent of prior knowledge of sequence variants. Window length shown is not on scale, typical window length for HiPlex sequencing is 80-150 bp.

**Figure 3.**
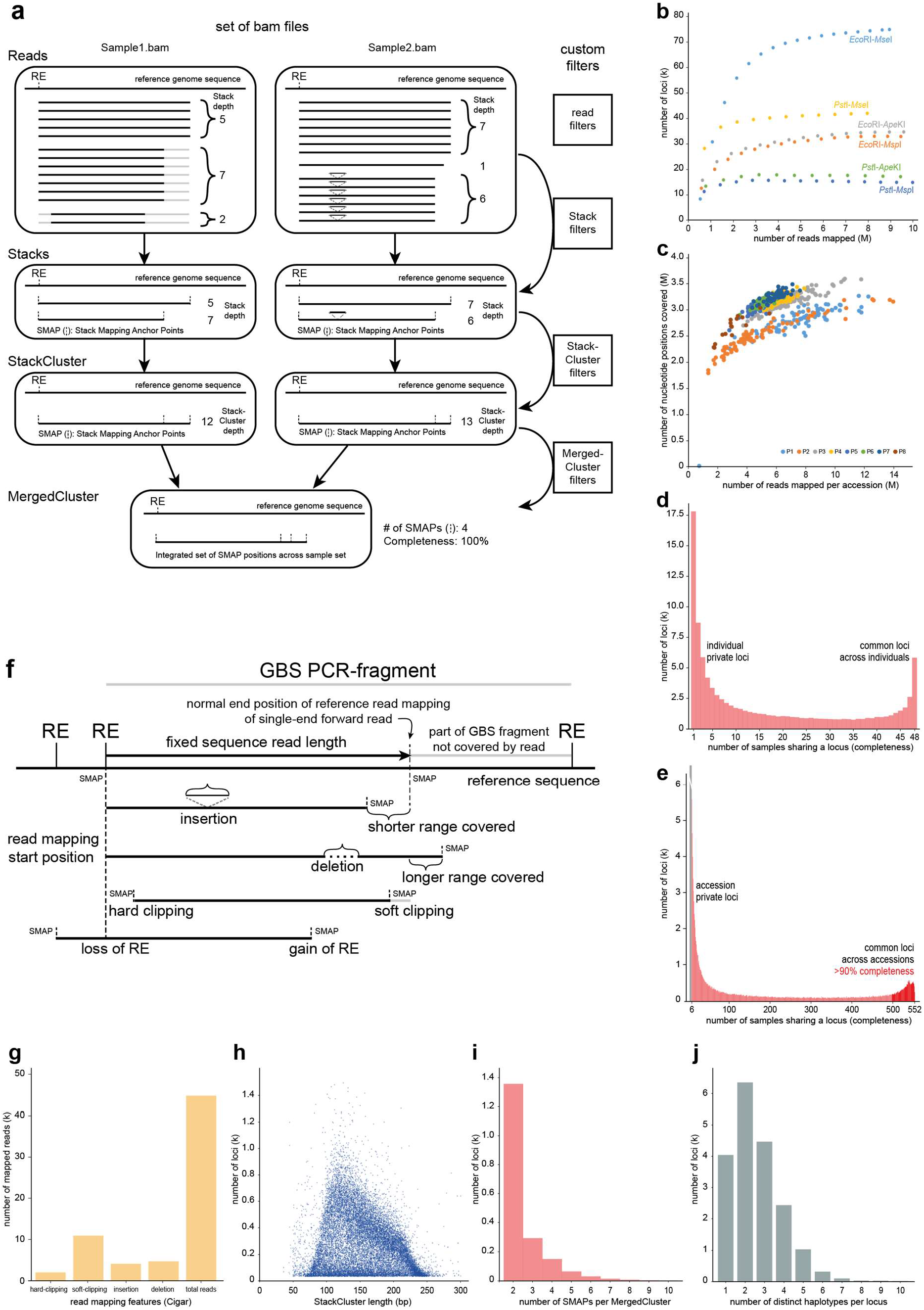
Scheme detailing the GBS locus delineation workflow of *SMAP delineate*. **a)** Module *SMAP delineate* analyses read distribution of reference-mapped GBS data to identify local read mapping position polymorphisms and to select regions of the genome with consistent read mapping across sample sets for downstream analysis such as variant calling and haplotype reconstruction. *SMAP delineate*’s working procedure includes delineating: (i) Stacks from reads with identical start and end mapping positions (SMAPs); (ii) StackClusters from overlapping Stacks within a sample; and (iii) MergedClusters by overlapping StackClusters across the entire sample set (**Supplemental Fig. S3**). *SMAP delineate* offers a range of key parameters to filter out SMAPs with low abundance relative to total locus read depth to control for noise in the data. *SMAP delineate* analyses GBS read mapping distributions per sample and plots the number of mapped reads against the number of detected loci, thus comparing: **b)** saturation curves of several enzyme combinations, *e.g.* to optimize GBS protocols in *M. sativa* (Julier *et al*., 2021), or **c)** to validate GBS Pool-Seq saturation across a collection of 768 *M. sativa* accessions divided in 8 subsets of 96 samples each (P1-P8); **d-e)** *SMAP delineate* plots the number of loci shared across a fraction of the sample set (completeness), *e.g.* for **d)** a set of 48 *L. perenne* individuals, or **e)** a set of 552 *L. perenne* accessions analysed via Pool-GBS (Blanco-Pastor *et al*., 2019, Keep *et al*., 2020); **f)** *SMAP delineate* further analyses read mapping polymorphisms within loci, and plots graphical summaries of: **g)** the abundance of partial alignments, **h)** read depth versus locus length distributions, and **i)** the distribution of the number of SMAPs per locus; **j)** Module *haplotype-sites* applied to *Pst*I-GBS of 48 *L. perenne* individuals shows the number of haplotypes per locus.

In brief, read data may be prepared for read mapping by our new tool *GBprocesS* (including demultiplexing, trimming, merging forward and reverse reads, quality filtering, etc.; see https://gbprocess.readthedocs.io/), and reads are mapped onto a reference sequence with any existing read mapper such as BWA-MEM (Li, 2013), Bowtie2 (Langmead *et al*., 2009), or Minimap2 (Li, 2018). Then, one of two SMAP modules may be invoked for the actual haplotyping. First, module *SMAP haplotype-sites* phases *a-priori* known ‘sites’ (defined as a series of polymorphic, single-nucleotide-resolution coordinate positions in the reference sequence), bundled at loci that are delineated depending on the type of sequencing library; *i.e.* based on primer positions for HiPlex data, sliding frames in Shotgun data, or, specifically for GBS data, Stack Mapping Anchor Points as identified with an additional tool called *SMAP delineate* (**Fig. 1** and **Supplementary Fig. S3**, and detailed explanation below). A set of indexed BAM files with mapped reads, a custom BED file with locus positions, and a VCF file with SNP or structural variant (SV) positions (identified by third-party tools such as GATK (Van der Auwera *et al*., 2020), Samtools (Li *et al*., 2009), or SNAPE-pooled (Raineri *et al*., 2012)), serve as input to *SMAP haplotype-sites*. Per read, the set of polymorphic sites are extracted from the read-reference aligned pair, and each site is scored as: absent “.”, reference “0”, alternative “1”, or gap “-”, thus transforming an aligned read into a read-reference-encoded, compressed haplotype string (**Fig. 1**).

The second module, called *SMAP haplotype-window*, delineates a locus by defining a so-called “window” enclosed by two locus-specific “border” sequences (coordinates are provided as a custom GFF file). *SMAP haplotype-window* then identifies the IDs of all reads mapped per locus in a BAM file; retrieves the corresponding sequences from the original FASTQ file (circumventing potential loss of sequence length due to soft-clipping or hard-clipping during read mapping); trims off the two locus-specific border sequences, and retains the entire remaining internal DNA sequence as a unique haplotype (**Fig. 2**). This concept for haplotyping is independent of *a-priori* variant calling, and is especially useful to *de novo* discover complex haplotypes derived from combined SNP and indel structures (*e.g.* derived from CRISPR/Cas genome editing, see De Bruyn *et al*., 2020).

Finally, in both modules, all unique haplotype counts are listed per locus per sample in an integrated haplotype call table. Both modules apply the same set of functionalities for haplotype frequency filtering, discrete genotype calling and locus and sample quality controls and create tabular and graphically represented summary statistics. The SMAP haplotype call table contains tabular formatted multi-allelic haplotype frequency spectra for Pool-Seq sample sets, or discrete multi-allelic haplotype calls for individual samples (diploids or polyploids), and scales from tens to thousands of loci and/or samples. A detailed comparison between *haplotype-sites* and *haplotype-window*, description of working procedure, guidelines for creating the required input files, utility tools, command line options, example data, and guidelines for troubleshooting are provided in the **Online Supplementary Material** and extensive **online manual** (see https://ngs-smap.readthedocs.io).

## Proof of concept and demonstration cases

We applied SMAP to a wide range of use cases and highlight specific functionalities per study. All studies have been described in full detail in the **Online Supplementary Material**.

First, *SMAP haplotype-sites* applied to a HiPlex panel of 339 multi-allelic regions sequenced in 765 autotetraploid potato lines, illustrates how SMAP haplotyping transforms the information content from around seven independent bi-allelic SNP markers per locus to a single multi-allelic marker with around six alleles per locus (**Supplementary Fig. S4a-b**). Furthermore, SMAP performs discrete haplotype dosage calling in autotetraploids (**Fig. 1g**); thereby enabling allele dosage per individual to be considered through a range of genetic models. Next, Shotgun targeted resequencing data obtained through probe capture, covering a total of 2.3 Mb genomic sequence across 503 candidate genes in 391 perennial ryegrass genotypes was analyzed with *SMAP haplotype-sites*, using dynamic sliding frames to bundle known neighboring SNP positions (Veeckman *et al*., 2019). This study revealed the influence of study-specific constraints on sample number, read length, library size (total number of reads sequenced per sample), SNP density, *within*- and *between*-sample genetic diversity etc., on a parameter optimization strategy balancing sliding frame length, minimal read depth, haplotype diversity per locus, and sample and locus call completeness (fraction of loci with minimal read depth per sample; fraction of samples with minimal read depth per locus) (**Supplementary Fig. S4e-h**). In addition, analysis of whole genome shotgun (WGS) sequencing with PacBio long reads in seven *Arabidopsis thaliana* ecotypes (Jiao and Schneeberger, 2020), revealed that those relationships hold true across sequencing platforms (**Supplementary Fig. S4i-l**), including long range sequencing technologies such as PacBio and, likely, Oxford Nanopore. Furthermore, exploiting the “semi-stacked” read mapping structure at the junctions of large scale deletions and inversions in WGS short read data of 269 *Oryza sativa* accessions (Kou *et al*., 2020), revealed how structural variants can be ‘haplotype’-encoded by scoring the absence/presence of read mapping in 3-bp frames at breakpoints (**Fig. 1a–c** panel Shotgun, locus 2: SV, and **Supplementary Fig. S5a-e**). Because of the nature of the SMAP haplotyping algorithm, SNPs that occur at any of the three interrogated nucleotides are called simultaneously and may create additional haplotype variants in the haplotype call table (using the “0” and “1” character types in the haplotype string). GBS is a popular method to sequence a representative fraction (typical range 0.1% - 1%) of a genome to generate genome-wide marker sets. GBS libraries are generated by ligation of sequencing adaptors to restriction enzyme overhangs, followed by size-selective PCR-amplification and sequencing (Elshire *et al*., 2011), *i.e.* the NGS equivalent of Amplified Fragment Length Polymorphism (AFLP). SNP calling algorithms typically rely on the comparison of ‘piles’ of aligned nucleotides to the respective reference nucleotide (**Supplementary Fig. S1**). So, constructing a complete genotype call table requires that a given nucleotide of the reference sequence is consistently covered by a minimal number of reads per sample across the sample set, and any factor that leads to missing read alignment creates missing data points in the genotype call table, thus lowering the power of GBS studies. We implemented *SMAP delineate* to systematically investigate GBS read mapping profiles prior to any variant calling (**Fig. 3a**), and disentangled the underlying technical and biological factors that cause variation in read mapping profiles. For instance, *SMAP delineate* saturation curves (**Fig. 3c**), reveal if the same *number* of loci are detected per sample (influenced by technical factors including the choice of enzymes and minimal required total library size (**Fig. 3b**); important for genetic studies that rely on linkage disequilibrium), while locus completeness plots reveal whether the *same* loci are detected across the sample set (**Fig. 3d,e**; *e.g.* influenced by biological factors such as the frequency of SNPs affecting restriction sites, **Supplementary Table S2**). *SMAP delineate* thus serves to routinely analyze read mapping distributions across multi-sample GBS datasets (**Fig. 3b–i**). Furthermore, we implemented the concept of Stack Mapping Anchor Points (SMAPs; **Fig. 3a,f**) to acknowledge the existence of *within*-locus read mapping polymorphisms (**Fig. 3g,i**), and integrated SMAPs with SNPs to create haplotypes with *SMAP haplotype-sites*, thus capturing read mapping polymorphisms as a novel type of molecular marker (**Supplementary Fig. S1**) and further enhancing the resolution of multi-allelic haplotypes in GBS data (**Fig. 3j**). Furthermore, a case study on the detection of interspecific hybrids in the *Festuca*-*Lolium* complex outlined how the SMAP toolset seamlessly iterates between untargeted molecular marker discovery (GBS), and targeted screening assays (HiPlex), and showed the added value of haplotyping for parental analyses and calculation of genetic similarity matrices (**Supplementary Fig. S6**). Finally, we demonstrated how *SMAP haplotype-window* characterizes multiplex CRISPR/Cas induced mutation spectra in two tetraploid potato cultivars, and showed how Pool-Seq combined with HiPlex genotyping facilitates highly efficient screening of large collections of mutants, while maintaining sensitivity for mutant allele detection (**Supplementary Fig. S7**).

### General conclusions and perspectives

The most prominent advantage of the SMAP package is that the haplotype table comprises a unified notation for multi-allelic variants that capture SNPs, SMAPs, SV junctions, and any combination thereof, so that all sequence variants per locus can be used together in a single integrated quantitative genetic analysis. We have highlighted the relative strengths and complementarity of novel concepts for haplotyping implemented in *SMAP haplotype-sites* and *haplotype-window*, and illustrated how various modules work together to extract the most informative haplotype diversity per sample set, depending on NGS library method, sequencing technology, prior knowledge of variants, and type of sample. For instance, initially conceived to explore the underlying causes of read mapping polymorphisms inherent to GBS, *SMAP delineate* now serves to routinely analyze read mapping distributions across multi-sample GBS datasets. Module *SMAP compare* shows the overlap in read mapping profiles across two sample sets (both analyzed by *SMAP delineate*), *e.g.* to optimize *SMAP delineate* parameters, to estimate reproducibility of GBS library preparation across laboratories, to compare GBS Pool-Seq to their constituent individuals, or to evaluate the number of common polymorphisms across genepools or breeding populations. The *SMAP utility* tools, in combination with complementary concepts to delineate loci such as sliding frames, allow to exploit many other types of NGS libraries and sequence polymorphisms, next to the most commonly used SNPs. Furthermore, *SMAP delineate* combined with *haplotype-sites* resolves the genotype calling paradox in GBS data by: 1) recognizing that read mapping polymorphisms exist; 2) systematically positioning read mapping polymorphisms in a coordinate system of SMAPs that mark locus start and end in a read data-driven fashion; and 3) combining SMAPs as a novel type of molecular marker with SNPs to create multi-allelic markers at the haplotype level. *SMAP delineate*, *haplotype-sites* and *haplotype-window* all offer a range of key parameters to filter features with low absolute and/or relative abundance at the level of SMAPs, SNPs, haplotypes, loci, and samples. Notably, parameters may be fine-tuned depending on the expected haplotype frequency, thus eliminating spurious signals and conferring robustness to discrete dosage haplotype calling in individuals, while facilitating sensitivity for detecting low frequency haplotypes in Pool-Seq data. Furthermore, analysis of WGS sequencing with PacBio long reads demonstrated that the SMAP haplotyping algorithm is scalable in all experimental dimensions (read length, numbers of loci, samples, and/or type of polymorphism), and that the application of SMAP is currently only limited by performance ceilings of sequencing technologies (including read length, read depth, and sequencing error rates).

We provide, and continue to develop, guidelines for optimization of SMAP parameter settings across a wide range of use cases in the **online manual**. All presented studies were based on Illumina short read data aligned with BWA-MEM (Li, 2013), or PacBio long reads aligned with Minimap2 (Li, 2018). As read-backed haplotyping is essentially dependent on the read-reference-pair alignment, users may test different read mappers in function of read length and/or genetic diversity between sample and reference. To avoid any potential coordinate shifts, *SMAP haplotype-sites* requires that the SNP variants are called on the same BAM files as used for haplotyping. We further envision that *SMAP haplotype-sites* can be used to quantitatively analyze RNA-Seq data for a variety of purposes using the haplotype count tables rather than the haplotype frequency tables: 1) to quantify expression of alternative splice-variants (analogous to SVs and using the genome sequence as reference for gapped alignments, *e.g.* **Fig. 1a–c**, Shotgun panel right locus); or 2) to quantify the relative expression level per haplotype based on sliding frames with neighboring SNPs to evaluate allele-specific expression bias using CDS transcripts as reference and avoiding gapped alignments. While simple haplotype frequency intervals worked surprisingly well to transform haplotype frequency spectra into discrete haplotype calls, we further see potential for using probabilistic models for inherent ploidy assignment and discrete genotype calling. Dominant and dosage calls are currently implemented, but all samples processed together need to be of the same ploidy level. To further develop the SMAP package, we are expanding the set of *SMAP utility* tools (**Supplementary Fig. S6**), to further support experimental design and downstream genetic analysis (*e.g.* a module called *SMAP design* that designs primers for HiPlex sequencing optionally combined with gRNAs for multiplex CRISPR/Cas genome editing). To facilitate downstream genetic analyses, *SMAP haplotype-sites* and *haplotype-window* optionally create the discrete haplotype call table in the format required for Cervus (Kalinowski *et al*., 2007), a program for parental analysis using multi-allelic genetic markers. Finally, we are developing a module called *SMAP effect-prediction* to exploit the benefits of sequence-based haplotypes generated by *SMAP haplotype-window* by reconstructing the genic context of the detected allelic sequence variants, translating the corresponding protein sequence, and quantitatively scoring the sequence similarity to the reference protein as proxy for the impact of an observed mutation on the protein functionality, in the frame of high-throughput screens for naturally occurring and/or CRISPR/Cas-induced mutations.

## Methods

Methods, including a detailed user manual explaining the features and optional parameters of the code, statements of data availability and any associated accession codes and references, are available in the **Online manual**, the **Online Supplementary Materials** and the **Online Supplementary Methods**.

## Data availability

SRA accession numbers of sequencing data that has been deposited in NCBI, are listed per study in the **Online Supplementary Methods**.

## Code availability

SMAP is available at https://gitlab.com/truttink/smap/ under the GNU Affero General Public License v3.0, and a detailed user manual is available at https://ngs-smap.readthedocs.io. Additional tools for downstream analysis of SMAP haplotype tables are available at https://gitlab.com/ybawin/smapapps and https://gitlab.com/ybawin/primer-design-gbs.

## Acknowledgments

We gratefully acknowledge Jens Renders, Rudolf Aelbrecht, and other students of the Ghent University 2019 Computational Biology class led by F.V. and P.D. for optimization of various parts of the initial code. We thank Hilde Muylle and Isabel Roldan-Ruiz for helpful discussions, Michaël Kelchtermans for his contributions to the online manual, and Marie Pégard for analyzing alfalfa data. This work was supported by the European Union’s Horizon 2020 research and innovation program (Marie Sklodowska-Curie Grant number 797162); the Irish Department of Agriculture, Food and the Marine (DAFM) through the Virtual Irish Centre for Crop Improvement (VICCI) (Project number 14/S/819); the Agency for the promotion of Innovation by Science and Technology (IWT) Flanders (project LO-080510); the Research Foundation Flanders (FWO Grant K208710N); the project GrassLandscape awarded by the 2014 FACCE-JPI ERA-NET+ call Climate Smart Agriculture (Grant Number: 609398) and by the EC (Grant number 618105), by the Agence Nationale de la Recherche (ANR) and the Institut National de la Recherche Agronomique (INRA - métaprogramme ACCAF) in France, the Biotechnology and Biological Sciences Research Council (BBSRC) in the United-Kingdom, the Bundesantalt für Landwirtschaft und Ernährung (BLE) in Germany; the European Union’s Horizon 2020 Program for Research & Innovation under grant agreement no. 727312 (project: “EUCLEG –Breeding forage and grain legumes to increase EU’s and China’s protein self-sufficiency”) and from the Ministry of Science and Technology of China (2017YFE0111000). The computational resources (Stevin Supercomputer Infrastructure) and services used for genotype calling and algorithm optimization were provided by the VSC (Flemish Supercomputer Center), funded by Ghent University in Belgium, FWO and the Flemish Government –department EWI. E.V. was supported by an ILVO PhD fellowship. A.L was supported by a joint PhD fellowship at ILVO and VIB (Vlaams Instituut voor Biotechnologie). Y.B. was funded by Research Foundation Flanders (FWO Grant G056517N).

## Author contributions

E.V. and T.R. conceived and designed the core of the SMAP code. D.S. designed and implemented GBprocesS and SMAP in all its current forms. Code optimization was performed by D.S., F.V and P.D. Test data analysis was performed by T.R., S.V.G., A.H., Y.B., J.L., P.B., S.Blugeon. M.L. and S.Byrne. T.R. wrote the manuscript with input from all authors. T.R. supervised development of the tool and guided the project to completion. All authors read and approved the final manuscript.

## ONLINE SUPPLEMENTARY MATERIAL

## ONLINE CONTENT

Any methods, additional references, source data, extended data, supplementary information, acknowledgments, details of author contributions and competing interests and statements of data and code availability, as well as a user manual with description of working procedure, guidelines for accurate read preprocessing and read mapping, command line options, use of arguments, and guidelines for troubleshooting, are available in the **Online Supplementary Material** and the **online manual** https://ngs-smap.readthedocs.io/.

**The Online Supplementary Materials provide the following parts:**

### Figures and tables

Figure S1 Difference in SNP versus haplotype calling and consequences for allele frequency estimation in Pool-Seq data.

Figure S2 The SMAP package seamlessly integrates subsequent modules into a workflow.

Figure S3 Scheme detailing haplotype definition in GBS loci by *SMAP delineate* and *haplotype-sites*.

Figure S4 *SMAP haplotype-sites* applied to HiPlex and Shotgun data.

Figure S5 Structural variant calling in *Oryza sativa*.

Figure S6 Detection of interspecific hybrids of *Festulolium*.

Figure S7 Multiplex CRISPR/Cas genome editing in potato.

Table S1 Non-exhaustive overview of various tools for haplotyping.

Table S2 Abundance of SNPs affecting restriction sites in *L. perenne*.

Table S3 Selection of 269 WGS datasets of *Oryza sativa* for haplotype calling of structural variants.

### Background information on the concepts

SNP calling versus haplotype calling.

Workflow ordering the components of the SMAP package.

Comparison between *haplotype-sites* and *haplotype-window*.

### Proof of concept and demonstration cases

### SMAP haplotype-sites

#### HiPlex

*The PotatoMASH genotyping tool applied to potato.*

#### Shotgun

*Sliding frames: haplotyping probe capture-enriched Shotgun data in perennial ryegrass.*

*Long reads: haplotyping of PacBio data WGS data in Arabidopsis.*

*Structural variants: haplotyping junctions of large scale deletions and inversions in WGS data in rice.*

### SMAP delineate

#### GBS

*Achieving saturated datasets: loci with sufficient data for genotype calling*

*Absence/presence of GBS loci due to SNPs at restriction sites*

*Absence/presence of read mapping at the nucleotide resolution: SMAPs*

*Using SMAPs as additional polymorphic marker*

### Iterative cycles of *SMAP delineate*, *SMAP haplotype-sites* and *SMAP utility* tools

#### GBS/HiPlex

*Identification of interspecific hybrids in the Festuca-Lolium complex*

### SMAP haplotype-window

#### HiPlex

*Multiplex CRISPR/Cas genome editing in potato*

**The Online Supplementary Methods provide the technical details of all demonstration cases.**

