## Supplementary material for "Stack Mapping Anchor Points (SMAP): a versatile suite of tools for read-backed haplotyping": Online_Supplemental_Materials

**The Online Supplementary Materials provide the following parts:**

**Figures and tables:**

Figure S1 | Difference in SNP versus haplotype calling and consequences for allele frequency estimation in Pool-Seq data.
Figure S2 | The SMAP package seamlessly integrates subsequent modules into a workflow.
Figure S3 | Scheme detailing haplotype definition in GBS loci by *SMAP delineate* and *haplotype-sites*.
Figure S4 | *SMAP haplotype-sites* applied to HiPlex and Shotgun data.
Figure S5 | Structural variant calling in *Oryza sativa*.
Figure S6 | Detection of interspecific hybrids of *Festulolium*.
Figure S7 | Multiplex CRISPR/Cas genome editing in potato.

Table S1 | Non-exhaustive overview of various tools for haplotyping.
Table S2 | Abundance of SNPs affecting restriction sites in *L. perenne*.
Table S3 | Selection of 269 WGS datasets of *Oryza sativa* for haplotype calling of structural variants.

**Background information on the concepts:**

SNP calling versus haplotype calling.
Workflow ordering the components of the SMAP package.
Comparison between *haplotype-sites* and *haplotype-window*.

**Proof of concept and demonstration cases**

***SMAP haplotype-sites***
  **HiPlex**
    *The PotatoMASH genotyping tool applied to potato.*
  **Shotgun**
    *Sliding frames: haplotyping probe capture-enriched Shotgun data in perennial ryegrass.*
    *Long reads: haplotyping of PacBio data WGS data in Arabidopsis.*
    *Structural variants: haplotyping junctions of large scale deletions and inversions in WGS data in rice.*

***SMAP delineate***
  **GBS**
    *Achieving saturated datasets: loci with sufficient data for genotype calling*
    *Absence/presence of GBS loci due to SNPs at restriction sites*
    *Absence/presence of read mapping at the nucleotide resolution: SMAPs*
    *Using SMAPs as additional polymorphic marker*

**Iterative cycles of *SMAP delineate*, *SMAP haplotype-sites* and *SMAP utility* tools**
  **GBS/HiPlex**
    *Identification of interspecific hybrids in the Festuca-Lolium complex*

***SMAP haplotype-window***
  **HiPlex**
    *Multiplex CRISPR/Cas genome editing in potato*

**Affiliations**
**Data availability**
**Code availability**
**References (continued)**

**The Online Supplementary Methods provide the technical details of all demonstration cases.**

**SNP calling versus haplotype calling**

**Supplementary Fig. S1** compares how allele frequency shifts between populations are calculated using SNPs or haplotypes. Because SNPs are shared between the different haplotypes, SNP allele frequency profiling (counting from top to bottom and ignoring the neighboring nucleotide context) estimates a reference allele frequency (RAF) shift from 55% to 30% RAF for the first SNP, from 40% to 50% RAF for the second SNP, and from 65% to 80% RAF for the third SNP, in population 1 versus population 2, respectively. Even if all chromosome equivalents have been taken into account (no false negative read depth observations), the reference allele frequency shifts are not strong (10-15%). The last SNP is characterized by partial loss of read alignment, so that the total read depth is higher in population 1 (121 counts) versus population 2 (66 counts). However, absolute read counts are typically not taken into consideration in population genetic analyses of allele frequency shifts, which focus on *relative* allele frequency shifts and consider read depth per nucleotide position in isolation (not noticing the local drop in read depth compared to neighboring nucleotides). The largest shift observed at the SNP level, is the right-hand side SNP that changes from 9% RAF in population 1 to 100% RAF in population 2. So, allele frequency shift analysis based on SNPs would identify the right-hand side SNP with the strongest signal, and likely disregard the allele frequency shifts at the other three SNPs. Haplotyping, according to *SMAP haplotype-sites*, looks at the locus from left-to-right at the read level, recognizes the partial absence of read mapping and encodes that signal into the haplotype string; separates the signals of SNPs that are shared by multiple haplotypes; counts each read per locus; and compares the frequencies for the four haplotypes. That test will show that haplotype “011.” increases relative frequency from population 1 (10%) to population 2 (50%), and the reference haplotype (“0000”) increases from 5% in population 1 to 30% in population 2. Consequently, the two other haplotypes must decrease proportionally as the total sum of relative frequencies is 100%, by definition. Haplotyping, therefore, presents a more accurate and realistic quantitative representation of genome equivalents and allelic diversity in Pool-Seq data compared to single SNP analyses.

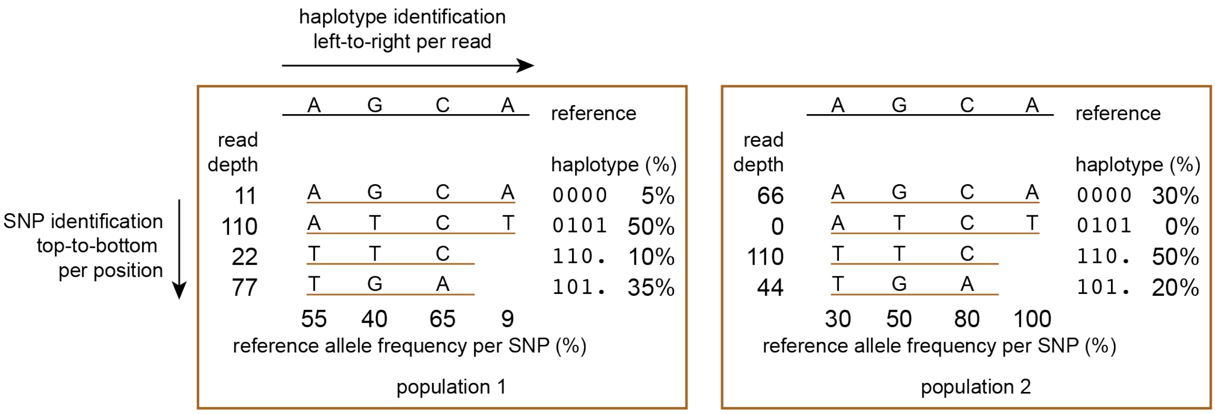


**Figure S1 | Difference in SNP versus haplotype calling and consequences for allele frequency estimation in Pool-Seq data.** Pool-Seq read data of two fictive populations are compared at a single locus. Both populations contain a set of haplotypes with different relative frequencies (for instance before and after selection for a beneficial allele, or against a detrimental allele). The observed read depths per haplotype per population are listed on the left-hand side of the stack. For ease of comparison, the total read depth per locus is the same. In ‘traditional’ SNP calling, read alignments are analyzed, nucleotide per nucleotide, independent of their sequence context (neighboring aligned nucleotides), and only positive observations (aligned nucleotides) are taken into consideration for SNP calling, as each pile of nucleotides is compared to the reference. In the sequence context that is covered by these GBS reads, four haplotypes exist that share at least three SNPs covered by all reads. In addition, a region exists in the reference genome that lacks read alignment (for any of the reasons outlined in **Fig. 3** and **Supplementary Fig. S3**) in two out of four haplotypes. This means that SNP calling algorithms are blind to the fact that genome equivalents exist (evidenced by the neighboring read alignment), but are not observed (false negative read depth at the right-hand SNP).

**SMAP package.** SMAP, including the source code written in Python is available at https://gitlab.com/truttink/smap/ under the GNU Affero General Public License v3.0. A detailed user manual describing the functionality of the modules of SMAP (**Supplementary Fig. S2**), and guidelines for accurate read processing is available at https://ngs-smap.readthedocs.io/. Tools for preprocessing of GBS reads are available at https://gbprocess.readthedocs.io/. Additional tools for downstream analysis of SMAP haplotype tables are available at https://gitlab.com/ybawin/smapapps and https://gitlab.com/ybawin/primer-design-gbs.

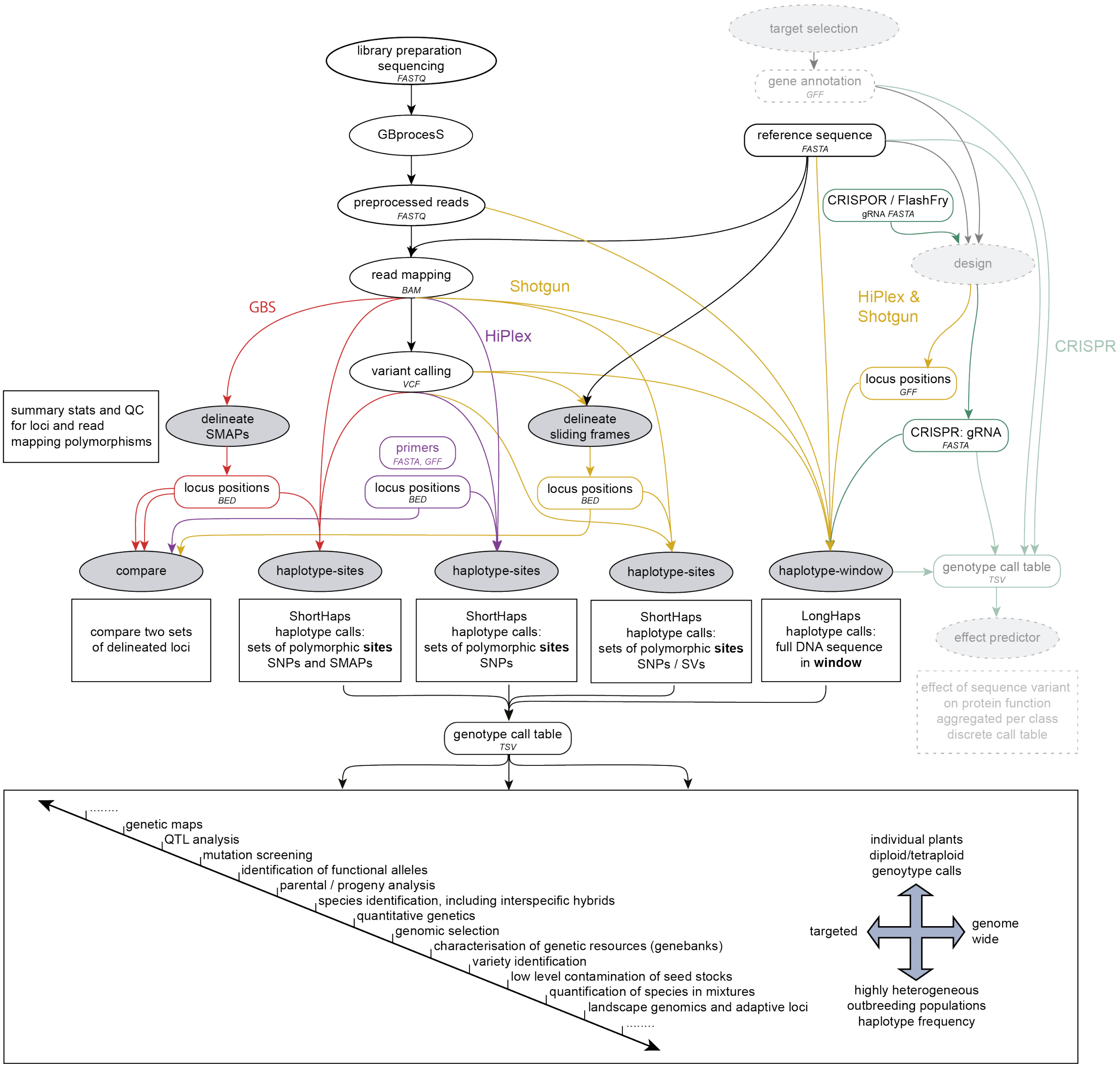


**Figure S2 | The SMAP package seamlessly integrates subsequent modules into a workflow**. The scheme displays a global overview of the functionalities of the SMAP package. White ovals are external operations and grey ovals are components of SMAP. Preprocessing of GBS reads should be performed by *GBprocesS* (see https://gbprocess.readthedocs.io/). Square boxes show the output of each of the components. Arrows show how output from various components are required input for the next component in the workflow for each of the NGS library types (GBS (red), HiPlex (purple), Shotgun (yellow)), file formats are shown in uppercase italics. Locus delineation is performed by *SMAP delineate* for GBS data, and sliding frames can be created with the *SMAP utility* tools. Haplotype calling is implemented in the components *SMAP haplotype-sites* and *haplotype-window*. Components *SMAP target-selection* (extracts and formats reference sequences and structural annotations of sets of candidate genes), *SMAP design* (designs multiplex amplicon primers (*e.g.* to screen for natural sequence variation in genepools, elite materials, or cultivars) and gRNAs for multiplex combinatorial CRISPR/Cas genome editing (green)) and *SMAP effect-prediction* (predicts the effect of sequence variants on the encoded protein sequence and functionality) are currently under development. Output from *SMAP haplotype-sites* and *SMAP haplotype-window* can be used for any downstream methods that use sets of targeted or genome-wide loci with multi-allelic markers, across the broad spectrum of genetic analysis, from QTLs on individuals up to landscape genomics on populations.

| software | algorithm | reference | prior knowledge | locus length | ploidy/Pool-Seq | imputation possible |
|---|---|---|---|---|---|---|
| TASSEL-UNEAK | reference-free identification of reciprocal tag pairs | reference-free | none | reads trimmed to 64 bp | any ploidy | no |
| Haplotag | read-backed haplotyping | reference-free | TASSEL-UNEAK output | reads trimmed to 64 bp | any ploidy | yes |
| GIBpPS | clustering of loci | reference-free | none | only tags with same length are clustered | any ploidy | no |
| fastPHASE | clustering of SNPs using a hidden Markov model that allows the cluster membership of observed haplotypes to change continuously along the genome | reference-free | SNP data and optionally known haplotypes | unlimited | diploids | yes |
| Hapler | read-backed consensus sequence construction minimizing possible crossovers | reference-based | SAM files and optional list of variant positions to consider | unlimited | unknown | no |
| LocHap-GBS | Locus windows are split into subwindows with maximum three heterozygous sites within any individual. Using the reads in each subwindow, LocHap-GBS uses a probabilistic model to identify haplotypes. | reference-based | bam files, bed file delineating the loci, VCF file with SNPs | max 3 neighboring SNPs | diploids | no |
| WhatsHap | read-backed or pedigree-based haplotype assembly using weighted minimum error correction | reference-based | VCF file with genotype calls per sample | phase extension of partially overlapping reads | any ploidy | no |
| harp | maximum likelihood estimation of allele frequencies using known haplotypes | reference-based | bam files, haplotypes should be known a priori | unlimited | any ploidy/pools | no |
| HapHunt BamBam | K-means clustering of loci | reference-based | bam files | unlimited | any ploidy | no |
| PoolHap | haplotype frequency estimation using a regression-based approach with known haplotypes | reference-based | bam files, haplotypes should be known a priori | unlimited | any ploidy/pools | no |
| ShoRAH | read-backed consensus sequence construction | reference-based | bam files | read length | any ploidy/pools | no |
| HaploJuice | estimation of the proportion of mixed subsamples using maximum likelihood followed by haplotype reconstruction using dynamic programming and selecting the best arrangement using maximum likelihood | reference-based | bam files, number of haplotypes | unlimited | any ploidy/pools | no |
| SMAP | read-backed haplotyping | reference-based | *haplotype-sites:* bam files, SNP, SV, SMAP coordinates<br>*haplotype-window:* bam files, fastq files, border coordinates | unlimited | any ploidy/pools | no |

| software | read data | link to software | literature reference | link to reference (doi) | remarks |
|---|---|---|---|---|---|
| TASSEL-UNEAK | short reads | https://www.maizegenetics.net/tassel | Glaubitz et al., 2014 | https://doi.org/10.1371/journal.pone.0090346 | TASSEL developers in 2015: "We are not spending any more time on UNEAK. I would really recommend making a very inexpensive pseudo reference, and just use it to anchor against and then run the GBSv2 pipeline." |
| Haplotag | short reads | depricated | Tinker et al., 2016 | https://doi.org/10.1534/g3.115.024596 | |
| GIBpPS | short reads | https://github.com/ahapke/gibpss | Hapke & Thiele, 2016 | https://doi.org/10.1111/1755-0998.12510 | |
| fastPHASE | short reads | http://scheet.org/software.html | Scheet & Stephens, 2006 | https://doi.org/10.1086/502802 | The models do not use the physical distances between markers. |
| Hapler | short reads | depricated | O'Neil and Emrich, 2011 | https://doi.org/10.1186/1471-2164-13-S2-S4 | |
| LocHap-GBS | short reads | http://compgenome.org/lochap/GBS/ | Manching et al., 2017 | https://doi.org/10.1534/g3.117.042036 | |
| WhatsHap | short and long reads | https://github.com/whatshap/whatshap | Patterson et al., 2015 | https://doi.org/10.1089/cmb.2014.0157 | Only suited for coverage up to 20x. It is recommended to use high quality short reads for variant calling and do the phasing with long reads. |
| harp | short reads | https://bitbucket.org/dkessner/harp/src/master/ | Kessner et al., 2013 | https://doi.org/10.1093/molbev/mst016 | |
| HapHunt BamBam | short reads | depricated | Page et al., 2014 | https://doi.org/10.1186/1756-0500-7-829 | |
| PoolHap | short reads | https://github.com/Gregor-Mendel-Institute/poolhap | Long et al., 2011 | https://doi.org/10.1371/journal.pone.0015292 | |
| ShoRAH | short reads | https://github.com/cbg-ethz/shorah | Zagordi et al., 2011 | https://doi.org/10.1186/1471-2105-12-119 | |
| HaploJuice | short reads | https://osf.io/b8nmf/ | Wong et al., 2018 | https://doi.org/10.1186/s12859-018-2424-7 | The method explores all the possible cases and is an exact algorithm, hence the execution time is growing exponentially with the number of haplotypes. It was designed and used for amplicon sequencing data. |
| SMAP | short and long reads | https://gitlab.com/truttink/smap/ | this publication | | *SMAP haplotype-sites* scores SNPs, SVs, and read mapping position polymorphisms<br>*SMAP haplotype-window* scores any sequence variant, independent of prior variant calling |

**Table S1 | Non-exhaustive overview of various tools for haplotyping.** For instance, UNEAK (Lu *et al*., 2013), GIBpPS (Hapke *et al*., 2016), and Haplotag (Tinker *et al*., 2016), each create haplotypes from GBS data using reference-free methods. Some algorithms focus on maximum likelihood estimation of allele frequencies based on known haplotypes (Kessner *et al*., 2013). LocHap-GBS (Manching *et al*., 2017) extracts local haplotypes based on a few (max 3) neighboring SNPs, and requires a predefined bed file that strictly delineates GBS tags using the restriction site distribution in the reference genome. WhatsHap (Patterson *et al*., 2015), performs phasing of a VCF file with predefined SNPs by extending the phase using locally overlapping reads.

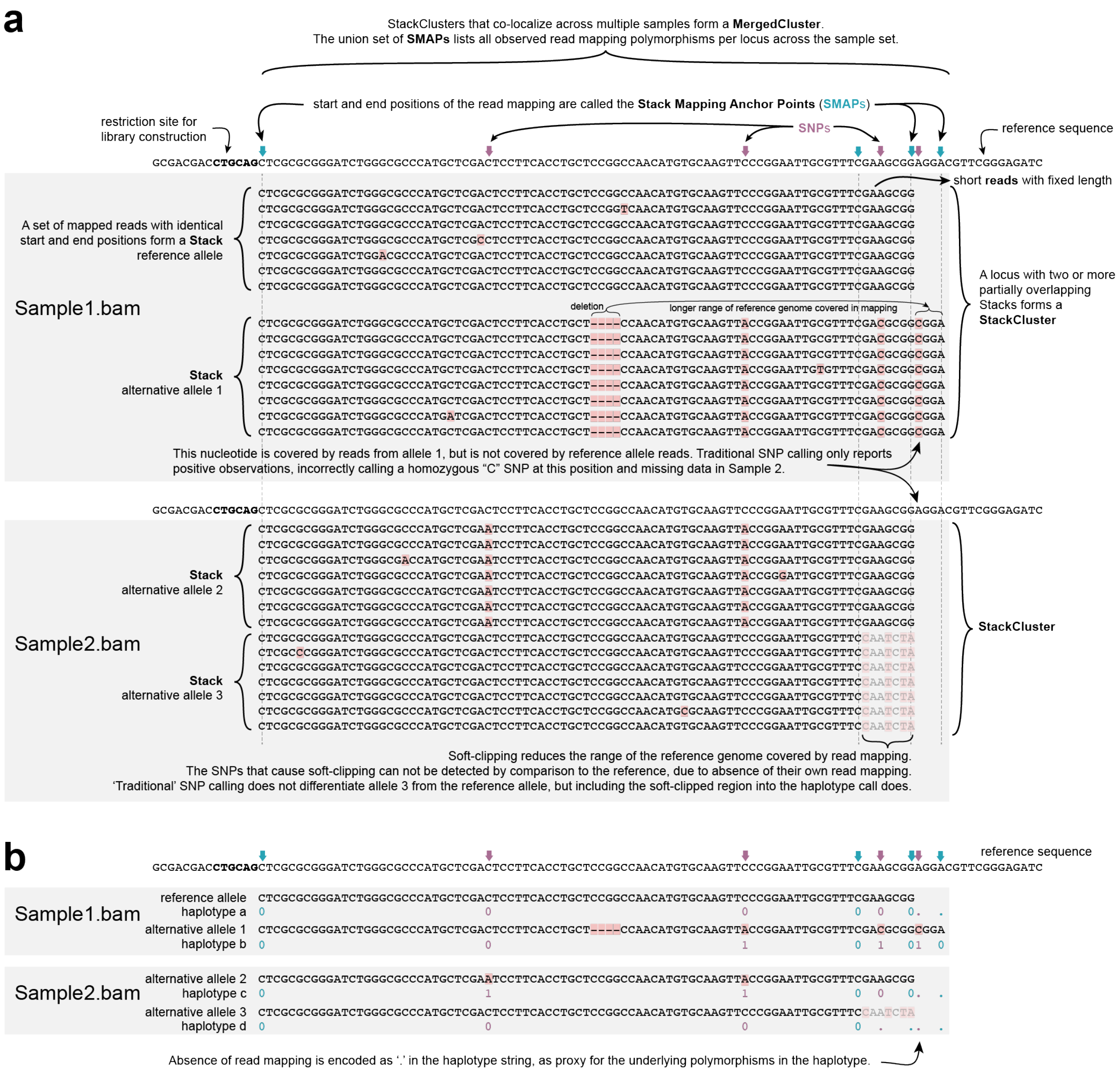


**Figure S3 | Scheme detailing haplotype definition in GBS loci by *SMAP delineate* and *haplotype-sites*. a)** definition of Stacks, StackClusters, and MergedClusters, and relative position of SMAPs and SNPs based on read-reference alignments in two heterozygous diploid samples displaying read mapping polymorphisms (SMAPs). Causes of read mapping polymorphisms and their consequences on SNP calling are indicated (see also **Fig. 3f**); **b)** scheme showing how a read-reference alignment is used to encode a compressed haplotype string by four character types: '0' for presence, reference nucleotide, '1' for presence, alternative nucleotide, '-' for gap in read alignment (not queried as no SNPs or SMAPs overlap with the deletion in allele 1), '.' for absence of read due to truncated read mapping. Note that while the length of the read mapping differs between alleles, all haplotype strings have the same number of characters, corresponding to the set of polymorphic sites (SMAPs and SNPs) in the locus. All non-polymorphic sites are skipped during haplotyping.

### Comparison between *haplotype-sites* and *haplotype-window*

While all SMAP modules pivot around the central theme of read-backed haplotyping, *haplotype-window* (**Fig. 2**) explores sequence variants that are out of reach from *haplotype-sites* (**Fig. 1**, **Fig. 3**). Here, we contrast the methods and illustrate their complementarity. First, *haplotype-sites* takes *a-priori* known sets of separate, single nucleotide positions, such as SNPs, SMAPs, and large scale inversion or deletion junctions as proxy for structural variants; scores the absence/presence and reference/alternative nucleotides in read-reference alignments, and aggregates neighboring sites per locus into a short haplotype string, while skipping internal, presumed non-polymorphic, sites. This yields the possibility to create multi-allelic haplotypes per locus, as long as more than 1 polymorphic site is included in the locus, and read mappings effectively span the length of the locus. In contrast, *haplotype-window* considers the entire native DNA strings as haplotype, without any required *a-priori* knowledge. Clearly, when prior knowledge of potentially polymorphic regions is available, this can be used to define custom sets of 'windows' enclosed by 'borders' at flanking non-polymorphic regions to target the analysis to those loci. While *haplotype-sites* considers SMAPs as the outer limits of loci and *includes* those characters in the haplotype string (**Fig. 3**), *haplotype-*

*window* sorts out reads based on global mapping to target loci, and searches for two border sequences within reads per locus, trims those off, and considers the entire DNA sequence in-between those border sequences as the haplotype (**Fig. 2**, *i.e.* the border sequences that delineate the locus are *excluded* from the haplotype string). While *haplotype-sites* strictly depends on the reference coordinate system of read-reference mappings and cigar strings, *haplotype-window* only uses the read mapping step to sort out reads across loci. The actual haplotypes are retrieved from the original FASTQ sequences (before read mapping) to circumvent sequence loss due to soft-clipping and hard-clipping. Where *haplotype-sites* strictly defines haplotype length per locus as the sum of all queried polymorphic sites, *haplotype-window* takes any sequence variant including any linear combination of insertions, deletions and/or SNPs, as a unique haplotype, of which the string length is defined by the observed sequence variants. This is especially important to discover complex yet local indel structures derived from naturally occurring sequence variants, or CRISPR/Cas or any other genome editing reagent, while circumventing specific indel detection algorithms that rely on read-reference read mapping on *both* sides of the indel. *Haplotype-sites* extracts only polymorphic sites from a broader genomic context, hence ignores sequence errors at internal (deemed non-polymorphic) positions; *haplotype-window* retains the entire sequence within a given locus as a full length haplotype, and is thus more sensitive to low-frequency sequencing errors that represent unique haplotypes. *Haplotype-sites* generates haplotypes encoded as "01.-" characters of fixed length per locus, while *haplotype-window* generates haplotypes as a string of ACGT characters (or IUPAC, *i.e.* the actual genomic DNA sequence) of any length. This last feature is especially important for the interpretation of the consequences of sequence variants on, for instance, encoded proteins, for which we are currently developing a novel module called *SMAP effect-prediction*. For example, CRISPR/Cas genome editing is widely applied to introduce mutations in target genes and the target loci can efficiently be re-sequenced using (HiPlex) amplicon sequencing to detect mutations. Most CRISPR-induced mutation detection software only scores mutations as sequence length difference to the reference (*e.g.*, +1 insertion; -4 deletion), but does not consider the exact position in the gene or open reading frame (ORF), nor its resulting effect on the actual translated protein (other than simply assuming that a non-triplet length difference leads to a frameshift mutation and knocks out the protein function). In contrast, *haplotype-window* extracts the observed haplotypes of the target loci, allows to locally substitute the original reference sequence with the newly observed allele, while accurately adjusting the coordinates of downstream intron-exon borders (in case of indels that change the reference sequence coordinate system), and is thereby able to predict the most likely translated protein given the adjusted gene model. This predicted protein is then compared to the reference protein by full length protein sequence alignment to quantitatively score the remaining sequence similarity as proxy for the impact of the observed mutation on the protein functionality.

**Proof of concept and demonstration cases**

In the following, we present different use cases to illustrate the application of *SMAP haplotype-sites* to HiPlex, Shotgun, and GBS datasets.

**HiPlex.** To demonstrate the principle of haplotyping in Illumina short read data, we applied *SMAP haplotype-sites* to HiPlex data. This dataset comprises amplicon sequencing of 765 autotetraploid potato lines using a genotyping tool called PotatoMASH (Potato Multi-Allele Scanning Haplotags), an amplicon panel covering 339 multi-allelic regions (165-180 bp) placed at 1 Mb intervals throughout the euchromatic portion of the potato genome. More details about the dataset and methods can be found below in the **Online Supplementary Methods**. We first detected 2,279 bi-allelic SNPs with a mean of 6.8 neighboring SNPs per locus. Next, we defined the start and end points for haplotypes (SMAPs) as the first and last nucleotide of the amplicon immediately internal to the primer pairs. Using *SMAP haplotype-sites* parameter settings for discrete dosage calling in tetraploid individuals (**Fig. 1a-c,** panel HiPlex). SMAP provides graphical output to confirm the expected haplotype frequency spectrum per sample given the organisms ploidy (**Fig. 1g**). The haplotype frequency spectrum is further used to transform the genotype call table into discrete haplotype dosage calls, after which loci with unexpected numbers of alleles per sample are removed from the dataset. This yielded 2,012 multi-allelic haplotypes in 334 loci corresponding to an average of 6 different short haplotypes at each locus across the population (**Supplementary Fig. S4a**). Thus, haplotyping transforms the information content per locus: from around seven independent bi-allelic SNP markers per locus to a single multi-allelic marker with around six alleles per locus. Finally, locus and sample call completeness and correctness scores (**Supplementary Fig. S4b-d**) are used to computationally filter low quality haplotypes, loci and/or samples from the final haplotype call table prior to performing any downstream genetic analysis. This use case demonstrates that SMAP can readily be applied to HiPlex data, particularly in outbreeding species where multiple variants occur within the short loci (80-200 bp) typically covered by amplicon sequencing. Furthermore, SMAP enables discrete haplotype dosage calling in autotetraploids; thereby enabling allele dosage to be considered through a range of genetic models.

**Shotgun.** Motivated by the advantages of customizing haplotype start and end points, we explored the possibility to perform haplotyping using dynamic sliding frames to bundle known neighboring SNP positions in Shotgun sequencing data, or, alternatively, to capture the junctions at large-scale deletions and inversions.

*Sliding frames*

If a read completely spans a given locus, an internal portion of that read can be used for read-backed phasing, and as long as a minimal read depth is available per locus, a reliable genotype call can be made per sample (**Fig. 1a-c,** panel Shotgun, locus 1: SNPs). Thus, known polymorphisms, like SNPs, can be grouped into customized sliding frames and define the start and end points for read-backed haplotyping. As a first proof of concept, we re-analyzed Shotgun targeted resequencing data (PE-91 reads) covering a total of 2.3 Mb genomic sequence across 503 candidate genes in 391 selected genotypes of *Lolium perenne* (Veeckman *et al*., 2019). Sliding frames were created with the *SMAP utility* tools, using the location of a given, known SNP as the locus start and the last SNP within a maximal distance (frame length) as the end point for haplotyping. Dynamically defined sliding frames thus create a typical distribution of locus length (**Supplementary Fig. S4e**) and number of SNPs per locus (**Supplementary Fig. S4f**), which follows the SNP density distribution along the chromosomes. The haplotype frequency spectrum revealed the highly heterozygous nature of the diploid samples (data not shown, a similar spectrum as **Fig. 1f**). This analysis further revealed the relationship between library size per sample (total number of reads sequenced), sliding frame length, and locus call completeness (fraction of loci with minimal read depth per sample) (**Supplementary Fig. S4g**). For instance, at increasing sliding frame length, less reads completely span the locus, reducing the total number of loci with minimum read depth. Conversely, increasing the library size can saturate locus call completeness per sample. An optimal sliding frame length can be identified for a given sequencing effort across the sample set with a trade-off between sliding frame length and sample call completeness. In addition, this analysis revealed that increasing the sliding frame length led to a higher average number of SNPs per locus (**Supplementary Fig. S4f**), and consequently a higher number of distinct haplotypes per locus, hence a higher information content per locus (**Supplementary Fig. S4h**). Depending on study-specific constraints on sample number, read length, library size, SNP density, *within*- and *between*-sample genetic diversity etc., users can perform a similar parameter optimization strategy balancing sliding frame length, minimal read depth, haplotype diversity per locus, and sample and locus call completeness.

*Long reads*

Next, we applied a sliding frame strategy to extract long range haplotypes from whole genome shotgun (WGS) sequencing with PacBio long reads (average read length around 7 kb) in seven *Arabidopsis thaliana* ecotypes (Jiao and Schneeberger, 2020). Sliding frames were created with 1.2M known SNPs (www.1001genomes.org) and the *SMAP utility* tools, scanning maximal frame length in the range 250-3000 bp (**Supplementary Fig. S4i**). The haplotype frequency distribution profiles of PacBio long read data display a relatively high proportion of spurious, low frequency haplotypes (**Supplementary Fig. S1e**), caused by a combination of relatively low read depth per locus, a high number of SNPs interrogated per locus (*e.g.* on average 10 SNPs and reaching up to 30 SNPs per haplotype in 1000 bp frames), and relatively higher sequencing error rates in PacBio long reads compared to Illumina short reads. In a predominantly homozygous organism such as *Arabidopsis*, such technical limitations can be mitigated by adjusting haplotype frequency interval boundaries for discrete genotype calling and selecting only loci with 100% sample call correctness (fraction of samples that show correct dosage score across the sample set, calculated per locus), across the sample set. Our analyses show that even with stringent parameter settings, a large number of loci (*e.g.* range 120k loci with 250 bp to 20k loci with 1000 bp frame length) with high quality haplotype calls can be extracted across the dataset (**Supplementary Fig. S4i-l**). These analyses further revealed that the relationships between sliding frame length, library size per sample (total number of reads sequenced), locus call completeness (fraction of loci with minimal read depth per sample), and extracted haplotype diversity per locus (**Supplementary Fig. S4i-l**), hold true across sequencing platforms, including long range sequencing technologies such as PacBio and, likely, Oxford Nanopore.

*Structural variants*

Next, we explored the possibility to genotype the breakpoints defining large-scale structural variants (SV), such as deletions (DEL) and inversions (INV) in WGS Illumina short read data of 269 selected *Oryza sativa* accessions of diverse origin, as previously reported by Kou *et al*., (2020). Here, haplotype calling exploits a "semi"-stacked read mapping structure. Read mappings start at randomly distributed positions (because of the nature of WGS libraries), but read alignment consistently stops at the site of the SV junction (because of the nature of read mapping algorithms), resulting in the sudden drop in read depth at the start and end points of deletions and inversions. This can be 'haplotype'-encoded by scoring the absence/presence of read mapping in a 3-bp frame: a central nucleotide at the breakpoint (defined in the structural variant VCF file), and its immediate nucleotide flanking positions (**Fig. 1a-c** panel Shotgun, locus 2: SV). Because of the nature of the *SMAP haplotype-sites* algorithm, SNPs that occur at any of the three interrogated nucleotides are called simultaneously and may create additional haplotype variants in the haplotype call table (using the "0" and "1" character types in the encoded haplotype string). The haplotype frequency spectrum revealed the highly homozygous nature of the diploid organism, with a haplotype frequency peak below 10% representing haplotypes derived from read errors (noise); the absence of a peak around 50% representing haplotypes at heterozygous loci; and a haplotype frequency peak at >90%, representing the homozygous, positively observed haplotype (**Fig. 1d**).

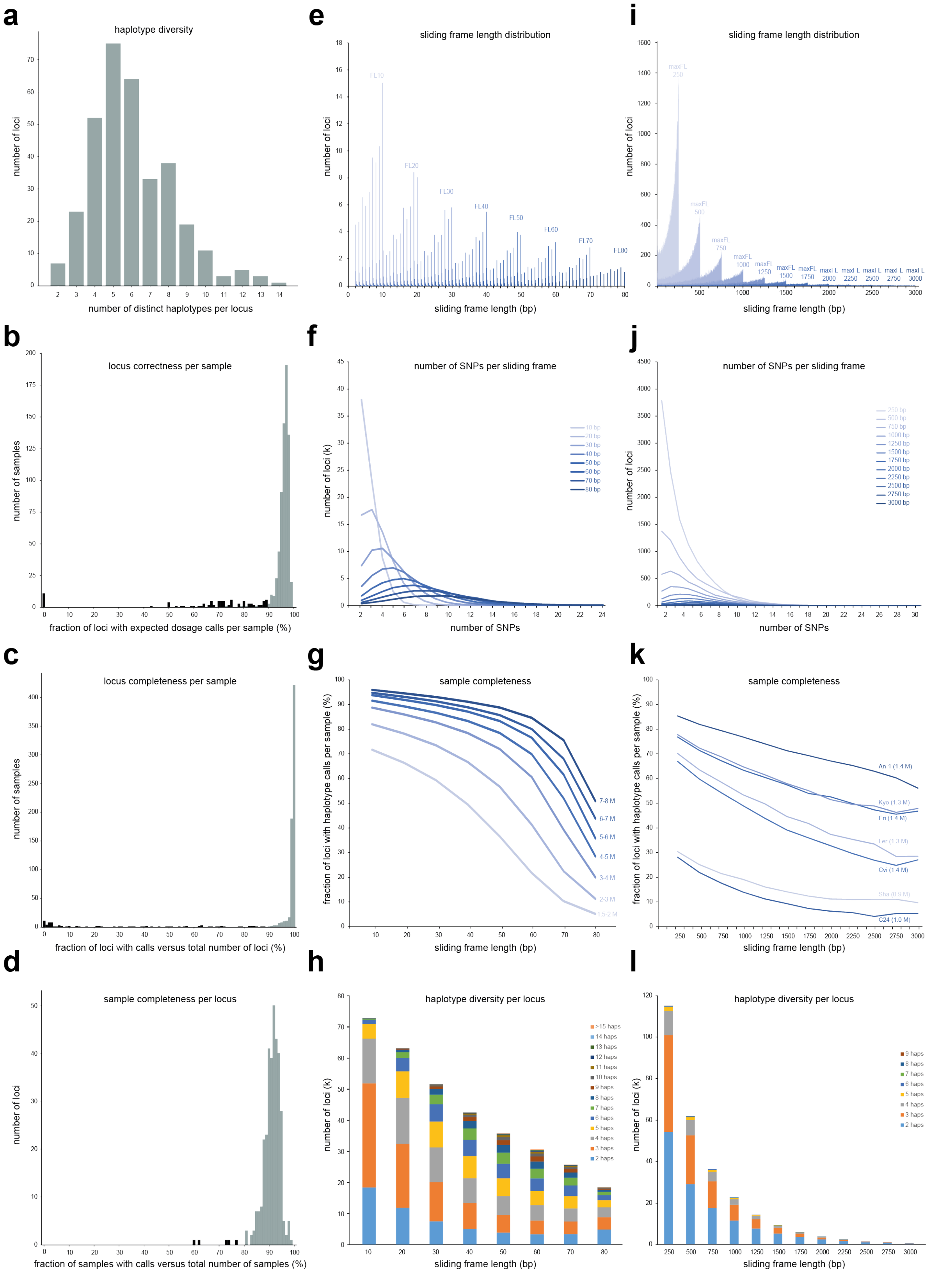


**Figure S4 | *SMAP haplotype-sites* applied to HiPlex and Shotgun data. a)** haplotype diversity (number of haplotypes per locus) of 334 loci of the PotatoMASH HiPlex amplicon panel across 765 *Solanum tuberosum* tetraploid individuals. **b)** locus call correctness; **c)** locus call completeness; and **d)** sample call completeness of the PotatoMASH HiPlex amplicon panel. Poor performing samples and/or loci (black bars, *e.g.* <90% locus correctness) may be identified and excluded from downstream analysis; **e-h)** haplotyping in sliding frames of Shotgun targeted resequencing data (PE-91 reads) covering a total of 2.3 Mb genomic sequence across 503 candidate genes in 391 selected genotypes of *Lolium perenne* (Veeckman *et al*., 2019). **e)** sliding frame length distribution at varying maximal sliding frame length (range 10-80 bp); **f)** distribution of the number of SNPs per locus at varying maximal sliding frame length (range 10-80 bp); **g)** sample call completeness at varying maximal sliding frame length.

Samples are grouped by library size class (total read count per library, range 1.5-2M to 7-8M) and completeness scores are averaged per group; **h)** haplotype diversity per locus, at varying sliding frame length. **i-l)** haplotyping in sliding frames of PacBio WGS data (average read length 7 kb) in seven *Arabidopsis* ecotypes (Jiao and Schneeberger, 2020); **i)** sliding frame length distribution at varying maximal sliding frame length (range 250-3000 bp); **j)** distribution of the number of SNPs per locus at varying maximal sliding frame length; **k)** sample call completeness at varying maximal sliding frame length; **l)** haplotype diversity per locus, at varying sliding frame length. Only loci with 100% locus call correctness across the seven samples are retained to extract high quality haplotype calls.

*SMAP haplotype-sites* can effectively be used to detect the SV junctions in neighboring deletions and large scale inversions as discrete dosage call in individuals with high locus completeness, provided that library size is sufficient to completely cover the genome at minimal read depth per locus (**Supplementary Fig. S5a**). The locus completeness score saturation curve constructed based on 71,973 known deletion junctions (Kou *et al*., 2020) on a set of 269 selected *Oryza* accessions (library size range 5 - 300M; PE-100 and PE-150 reads), shows the relationship between read length, library size, and minimal read depth, and indicates that around 50M PE-150 WGS reads are required for saturating genotype calling at minimal read depth of 15 or 200M PE-150 WGS reads at minimal read depth of 30 (**Supplementary Fig. S5b**). Similar results were obtained scoring a random selection of 99,886 known Inversion junctions in the same samples (**Supplementary Fig. S5d**). As a summary statistic, *SMAP haplotype-sites* calculates the relative population allele frequency across *Oryza sativa* accessions *per* haplotype (*i.e.* the abundance per junction variant within the species). Analysis on a subset of 100 accessions (with sufficient locus completeness) shows that, as expected, the reference haplotype (encoded as "000") is the major allele (range 80-100% haplotype frequency in the sample collection) across virtually all loci tested, and the observed structural variant alleles show minor haplotype frequencies across the population (range 1-20%). A similar fraction of "SNP-only" sequence variants is additionally identified at the same 3-bp locations in genotypes that do not contain a structural variant junction but incidentally contain SNPs at those loci (**Supplementary Fig. S5c** and **Fig. S5e**), thus further increasing haplotype diversity information content per locus.

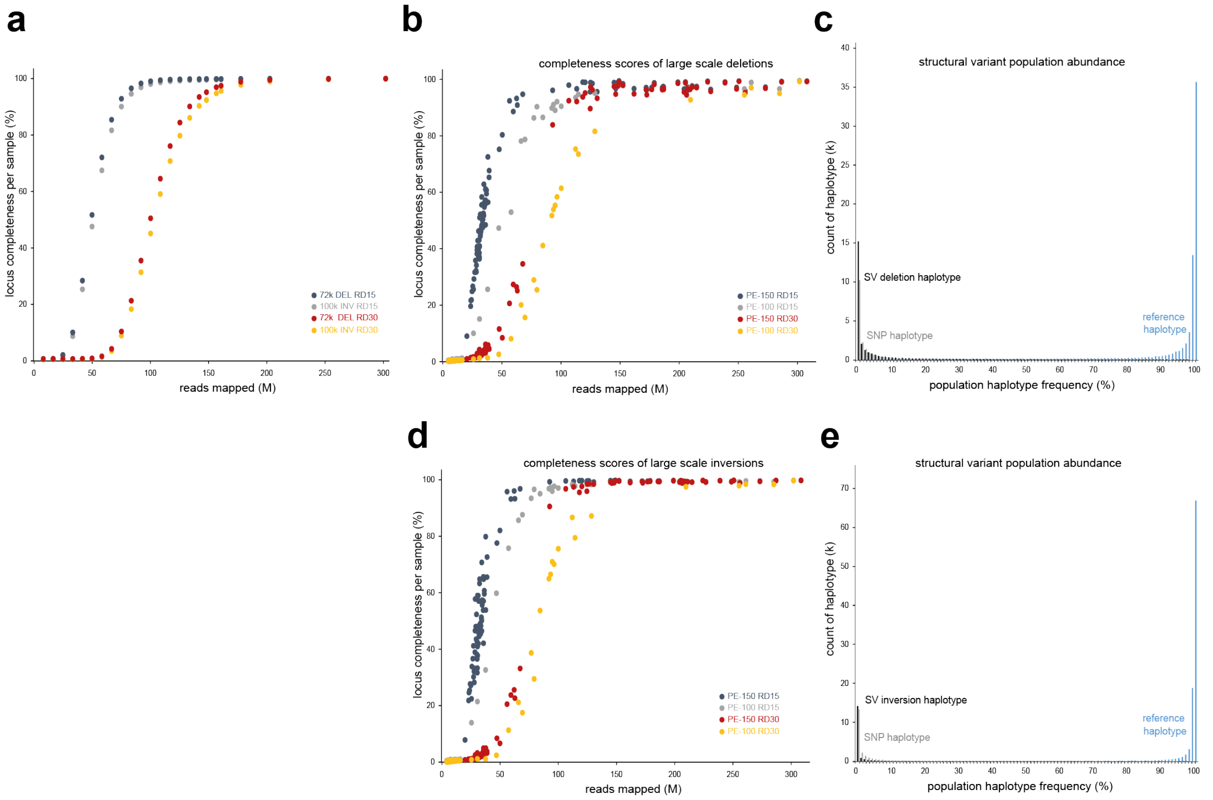


**Figure S5 | Structural variant calling in *Oryza sativa*. a)** Saturation curve for 72k DEL and 100k INV structural variants called on Nipponbare WGS data at increasing numbers of reads mapped. Non-saturated samples are created by computational subsampling from a fastq file with maximum 301M reads. Locus completeness is scored per sample at minimum read depth of 15 (RD15) or 30 (RD30). **b)** locus completeness score saturation curve constructed for 71,973 known deletion junctions (Kou *et al*., 2020) on 269 selected *Oryza sativa* accessions (library size range 5 - 300M; PE-100 and PE-150 reads), at minimal read depth per locus of 15 (RD15) or 30 reads (RD30). **c)** relative haplotype frequency across 100 *Oryza sativa* accessions (*i.e.* deletion junction variant population abundance). **d)** locus completeness score saturation curve constructed for 99,886 known inversion junctions. **e)** relative haplotype frequency across 100 *Oryza sativa* accessions (*i.e.* large scale inversion junction variant population abundance). The reference haplotype (blue bars) is the major allele (right-skewed distribution) across virtually all loci tested. Alternative haplotypes are aggregated by structural variants (SV, junctions of large scale inversions or deletions, black bars) or 'SNP-only' haplotypes (grey bars). All alternative haplotypes display a left-skewed population abundance distribution.

**GBS.** GBS is a type of reduced representation library generated by ligation of sequencing adaptors to restriction enzyme overhangs, followed by size-selective PCR-amplification and sequencing (Elshire *et al*., 2011), *i.e.* the NGS equivalent of Amplified Fragment Length Polymorphism (AFLP).

*Achieving saturated datasets: loci with sufficient data for genotype calling*

Before genotyping a large set of samples with GBS in a novel species with a given genome size and SNP density, it is essential to optimize the combination of restriction enzymes (one or two enzymes and their cutting frequency) and size selection stringency in order to yield the minimally required number of loci for the study (*i.e.* genetic resolution). Furthermore, mapping a minimal number of reads for each locus is needed for accurate genotype calling, and thus requires a saturating sequencing effort per sample. However, unequal distribution of library size across the sample set, combined with overall non-saturating library size often lead to insufficient read depth at increasing numbers of loci and/or samples in GBS studies, hence, missing data in the genotype call table.
To support library preparation method optimization, module *SMAP delineate* analyses GBS read mapping distributions per sample (**Fig. 3a**) and plots the number of mapped reads against the number of loci, thus creating comparative saturation curves that indicate the expected fraction of missing data per sample at a given total library size. For example, from a set of six enzyme combinations tested in a pilot experiment, *Pst*I-*Mse*I double-enzyme GBS performed on *Medicago sativa* (**Fig. 3b**) yields a plateau of 10k GBS loci with 10M mapped reads per sample (Julier *et al*., 2021). Given the optimal RE combination and known minimal required number of reads per sample, a set of 1061 samples was then genotyped with the appropriate sequencing effort. Subsequently, *SMAP delineate* was used to confirm locus saturation per sample across the sample subsets and/or to calibrate additional sequencing effort for drop-out samples (**Fig. 3c**). Following this scenario, we could generate 228,568 SNP markers on 31,743 loci with less than 5% missing locus-sample data points on 1061 *M. sativa* accessions. For genetic studies that rely on LD, it is important to estimate the distance between loci; in this case "stacks" of GBS reads. More specifically, to distinguish: 1) *between*-locus distance (genome size divided by number of GBS loci with sufficient data); from 2) the distance *between* neighboring loci with actual polymorphisms; and from 3) *within*-locus SNP distance (total number of SNPs divided by total length of all sequenced loci). *SMAP delineate* is run independent of SNP calling and specifically reports the number of detected GBS loci, providing a measure for *between*-locus distance. In line with this distinction, the subsequent module *SMAP haplotype-sites* (see below) can be used to analyse the number of loci actually containing SNPs and, as derivatives, the distance *between* loci with polymorphisms and the *within*-locus SNP distance.

*Absence/presence of GBS loci due to SNPs at restriction sites*

*SMAP delineate* can further be used to estimate to what extent samples contain not only the same *number* of loci (*i.e.* library size saturation), but also share the *same* loci (required to create a complete genotype table with SNP observations of all samples per locus) by analysing positional overlap of detected loci across the sample set (**Fig. 3a**). This analysis is driven by the notion that SNPs may cause the gain and loss of restriction sites, hence, cause the presence/absence of GBS loci in a genotype-dependent manner, inflating the total number of loci observed at least once across the sample set, while reducing the number of *shared* loci across all samples. Therefore, we analysed a set of previously identified SNPs based on targeted resequencing in a highly diverse outbreeding species *L. perenne* (Veeckman *et al*., 2019), and found that a substantial portion of SNPs co-localised at potential restriction sites and displayed variable frequency across the collection of 736 individuals (**Supplementary Table S2**). A high level of genetic diversity is expected to lead to a high fraction of loci that are either unique to, or lost from, a subset of genotypes. The locus completeness plot of *SMAP delineate* reveals the number of samples that share a given locus, *e.g.*, across a set of 48 *L. perenne* individuals (**Fig. 3d**). Absence of read mapping thus eliminates the possibility to call SNPs at those loci for a fraction of the samples, and creates missing data in the genotype call table that can never be mitigated by additional sequencing depth. Likewise, Pool-Seq data of 552 natural accessions of *L. perenne* (Blanco-Pastor *et al*., 2019, Keep *et al*., 2020) shows the high abundance of accession-private loci (**Fig. 3e**), in line with the high degree of genetic diversity across the species' natural range. Note that the analysis was performed with minimum completeness at 1%, meaning that 138,513 loci are shown that were observed in at least 6 out of 552 accessions. An additional 111,998 unique loci were observed only in 1 up to 5 accessions (grey bars in **Fig. 3e,** of which 56,143 strictly "single accession"-private, but are excluded because they have no data in any other accession. Conveniently, *SMAP delineate* provides the tools to select only the common loci shared across the majority of accessions (*e.g.* 19,487 loci common in at least 497 accessions, >90% sample completeness, red bars in **Fig. 3e**), prior to SNP calling and further genetic analysis.

| enzyme | effect | SNPs MAF>5% count | (%) | MAF1%-5% count | (%) | MAF<1% count | (%) | all SNPs count | (%) | number of RE in reference genome (2.3 Mb) |
|---|---|---|---|---|---|---|---|---|---|---|
| *Pst*I | loss | 177 | (14.5) | 103 | (8.5) | 435 | (35.7) | 715 | (58.7) | 1218 |
| | gain | 129 | (10.6) | 95 | (7.8) | 316 | (25.9) | 540 | (44.3) | |
| | shift | . | . | . | . | 1 | (0.1) | 1 | (0.1) | |
| *Eco*RI | loss | 90 | (15.0) | 57 | (9.5) | 182 | (30.3) | 329 | (54.7) | 601 |
| | gain | 82 | (13.6) | 67 | (11.1) | 245 | (40.8) | 394 | (65.6) | |
| | shift | . | . | . | . | . | . | . | . | |
| *Msp*I | loss | 816 | (9.4) | 648 | (7.5) | 2235 | (25.7) | 3699 | (42.6) | 8690 |
| | gain | 677 | (7.8) | 457 | (5.3) | 1506 | (17.3) | 2640 | (30.4) | |
| | shift | 19 | (0.2) | 10 | (0.1) | 38 | (0.4) | 67 | (0.8) | |
| *Mse*I | loss | 889 | (8.2) | 535 | (4.9) | 1987 | (18.4) | 3411 | (31.5) | 10820 |
| | gain | 834 | (7.7) | 683 | (6.3) | 2402 | (22.2) | 3919 | (36.2) | |
| | shift | 79 | (0.7) | 52 | (0.5) | 145 | (1.3) | 276 | (2.6) | |

**Table S2 | Abundance of SNPs affecting restriction sites in *L. perenne*.** | The relative proportion of restriction sites affected by SNPs is presented as percentage of the total number of restriction sites in the sequenced genomic regions.

*Absence/presence of read mapping at the nucleotide resolution: read mapping position polymorphisms*

The common intuitive expectation of reference mapped GBS reads is that all reads derived from a given locus have the exact same mapping start and end positions (commonly known as "stacks" of reads). This is because in principle, GBS reads are "anchored" to restriction sites, sequenced with fixed length (typically Illumina SE or PE short reads) followed by seeded alignment extension during read mapping (*e.g.* BWA-MEM). Inspired by the notion that 'classical' AFLP displays genetic polymorphisms as absence/presence of fragments of different length separated by gel electrophoresis, we set out to explore if mapped GBS reads similarly showed "read mapping position polymorphisms" at the nucleotide level. We expanded on the concept of "Stacks" by introducing StackClusters and MergedClusters, all delineated by so-called Stack Mapping Anchor Points (SMAPs), defined as the alternative read mapping start and end points at the same locus, due to insertions, deletions, soft-clipping and/or hard-clipping at the read mapping level (**Fig. 3f**, **Supplementary Fig. S3**). We implemented these concepts in *SMAP delineate* and applied it to a broad range of species (*L. perenne*, *Coffea arabica*, *Trifolium pratense*, *M. sativa* and *Rosa wichuriana*, see **online manual**), sample types (diploids, tetraploids, and Pool-GBS), and GBS library variants: single-digest GBS or double-digest GBS; combined with single-end or paired-end reads; separately mapped or merged (Zhang *et al*., 2014) paired-end reads before mapping.

Taken together, our comprehensive GBS read mapping analyses exposed a paradox underlying SNP calling in GBS data (**Fig. 3f-j**, **Supplementary Fig. S3**). While GBS reads mapped onto a reference genome sequence are used to identify SNPs, at the *within*-locus (nucleotide) level indels and SNPs (indirectly leading to soft-clipping or hard-clipping) themselves define where reads do and do not map onto the reference genome sequence. Because read-reference alignment is required to call a polymorphism, SNPs and indels thus hamper their own detection or that of neighboring polymorphisms in a given haplotype context (**Supplementary Fig. S3**). Furthermore, read-backed haplotyping requires the definition of the start and end point of loci to bundle sets of polymorphic 'sites' into a string of consecutive phased markers. Again, not only restriction sites and read length (which may theoretically be used to *in silico* predict locus coordinates on the reference genome sequence), but also SNPs and indels themselves define which regions on the reference genome sequence are actually covered by reads that diverge from the reference sequence (**Fig. 3f**), and thus influence the start and end point of their own loci. *SMAP delineate* provides tools to estimate to which extent this occurs in any given GBS dataset. It creates graphical summaries of the abundance of reads with insertions, deletions, soft-clipping, and hard-clipping in each BAM file (**Fig. 3g**); of read depth and read length distributions (**Fig. 3h**), and of the number of alternative SMAPs per locus, for all loci per sample and across all samples per locus (*i.e.* MergedClusters) (**Fig. 3i**). For instance, *Pst*I single-digest GBS data of 48 diploid *L. perenne* individuals reveals that at least one-third of all loci display *within*-locus read mapping polymorphisms (**Fig. 3i**).

*Using SMAPs as additional polymorphic marker*

Because SNPs and indels cause the alternative read mapping patterns per locus, the presence/absence of read mapping at Stack Mapping Anchor Points (SMAPs) reflect true polymorphisms. *SMAP haplotype-sites* has the ability to score the absence/presence of read mapping at SMAPs as a new type of polymorphism in addition to SNPs, and integrate all in a single type of haplotype string (**Fig. 1c** panel GBS, **Fig. 3** and **Supplementary Fig. S3**). Here, haplotype strings are encoded by 4 characters: '0' for presence, reference nucleotide, '1' for presence, alternative nucleotide, '-' for gap in read alignment, '.' for absence of read (i.e. due to truncated read mapping). Interrogated sites per loci are the outer SMAPs, any internal SMAPs (defined by *SMAP delineate*), and any locus-overlapping SNPs (defined by third-party SNP calling software). **Supplementary Fig. S3** illustrates how *SMAP haplotype-sites* uses SMAPs and SNPs to define haplotype strings per GBS read. The haplotype encoding takes the information derived from 'truncated' read mapping as a proxy for polymorphisms that were not (indels) or cannot (soft-clipping, hard-clipping) be detected due to -paradoxically- the same absence of read mapping. For instance, *Pst*I single-digest GBS data of 48 diploid *L. perenne* individuals shows a high number of distinct haplotypes per locus, thus increasing the genetic resolution of GBS marker loci (**Fig. 3j**). *SMAP haplotype-sites* was further tested on real sequencing data of highly heterozygous diploid and tetraploid individuals and pooled populations (Pool-Seq) of a wide range of species including *L. perenne*, *Coffea arabica*, *Trifolium pratense*, *M. sativa*, and *Rosa wichuriana* (see **online manual**).

**Iterative cycles of SMAP: identification of interspecific hybrids in the *festulolium* complex.** Next, we demonstrate how iterative rounds of SMAP haplotyping can be used to fine-tune genetic analyses. With the purpose to create interspecific hybrids within the *Festuca-Lolium* complex, we selected 11 *Lolium perenne*, 12 *Lolium multiflorum*, and 14 *Festuca pratensis* parental lines from a breeding program. *Pst*I-GBS (Verwimp *et al*., 2018), was performed on each individual to estimate genetic diversity *within* and *between* the species using *SMAP delineate* and *SMAP haplotype-sites* for haplotype calling. Next, *SMAP utility* tools were used to create a pairwise genetic similarity matrix and to identify discriminatory loci (loci without any shared haplotypes, *e.g.* 'a' vs 'bc', or 'ad' vs 'ef') between each pairwise comparison. HiPlex primer design *within* GBS loci taking all known SNPs into account created a set of 272 HiPlex amplicons, spread across the seven chromosomes of *L. perenne* (Nagy *et al.*, submitted for publication), that together yielded discriminatory power between all pairwise genotype combinations (**Supplementary Fig. S6**). Thus, a single HiPlex PCR per sample makes it possible to detect the presence of *L. perenne*-private, *L. multiflorum*-private and *F. pratensis*-private haplotypes. In parallel, manual interspecific crosses were performed between *L. multiflorum* and *F. pratensis*. Embryo rescue yielded 88 seedlings that were genotyped together with the parental lines with the HiPlex assay. Out of 85 progeny plants sequenced, 38 hybrid seedlings were identified and were distinguished from 47 non-hybrid seedlings (**Supplementary Fig. S6**). The genetic similarity matrix shows the versatile use of SMAP haplotyping. First, it shows the *within* and *between* genetic similarity of the selected parental lines and species, in which *F. pratensis* shows the least genetic diversity (94-100% loci with common haplotypes), and *L. multiflorum* the highest genetic diversity (81-96% loci with common haplotypes). Interspecific hybrids are identified because they share common alleles *both* with all *L. multiflorum* individuals, and display near-100% similarity with their actual *L. multiflorum* parent, *and* they share common alleles with all *F. pratensis* individuals and again near-100% similarity with their actual *F. pratensis* parent. The identity of the non-hybrid seedlings (absence of high levels of *F. pratensis* common alleles), is resolved by the marked high shared similarity with the *L. multiflorum* mother plant (as expected) and one alternative other *L. multiflorum* line of the same set. Thus, a purposefully designed set of 272 multi-allelic markers provides sufficient resolution for parental analysis both within species and across species in interspecific hybrids.

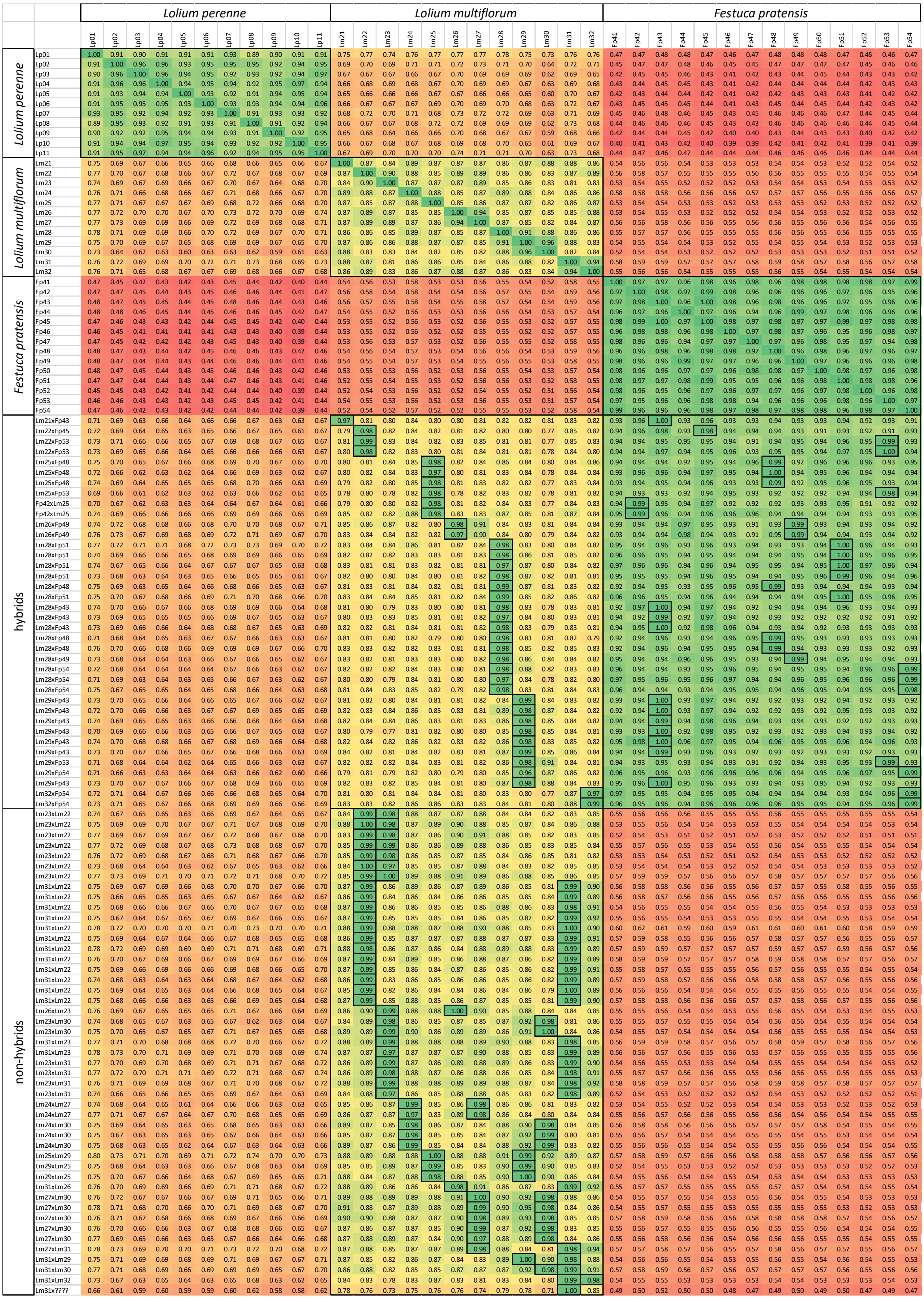


**Figure S6 | Detection of interspecific hybrids of *Festulolium*.** Genetic similarity matrix of the selected parental lines and potential hybrids. Each row/column represents a plant. 272 polymorphic HiPlex markers were developed that together discriminate 11 *Lolium perenne*, 12 *Lolium multiflorum* and 14 *Festuca pratensis* parental lines, based on shared/unique haplotypes. Out of 85 progeny plants screened, 38 *L. multiflorum* x *F. pratensis* hybrid seedlings were identified and were distinguished from 47 non-hybrid seedlings identified as progeny from two *L. multiflorum* lines. The *SMAP utility* tool *SMAPapp-Matrix.py* calculates the percentage of loci with partially shared alleles. The highest percentage corresponds with the respective parents.

**Multiplex CRISPR/Cas genome editing in potato.** A case study illustrates the analysis of multiplex CRISPR/Cas genome editing in potato (**Supplementary Fig. S7**). Protoplasts of 'Bintje' and 'Spunta' cultivars were co-transfected with a Cas9 expression vector and sets of separate expression vectors each containing a single gRNA using using polyethylene glycol (PEG). Either small pools of 11 gRNA vectors or large pools (22 gRNA vectors of two small pools combined) were used for co-transfection. Regenerated calli were genotyped by a single HiPlex PCR to simultaneously resequence all targeted loci. *SMAP haplotype-window* revealed different dosage per allele (*e.g.* loci EDR2_3, PSY1R_3, BON3_3, WAT1_3), different sets of alleles (*e.g.* loci BON1_1, PSKR1_1, FER_3, DMR_1), in the tetraploid Bintje and Spunta non-transfected control materials (compare samples S01 and S16), and some alleles with a SNP in the gRNA/PAM target sequence (*e.g.* PSKR1_1 and FER_3 in Spunta). Pooling multiple calli in one DNA extract (in this study five calli per pool) facilitates efficient screening of large collections of transfected materials, while maintaining sensitivity for mutant allele detection (a haplotype frequency of 5% corresponds to the detection of a single allele edited in a single callus in a pool of five). Screening of single calli can quantitatively highlight which alleles (out of four) have been edited and which alleles remained identical to their wild type (wt) reference sequence. Some gRNAs only led to edits in single samples (*e.g.* BON1_1), while other gRNAs (*e.g.* targeting locus EXLA2_2) induced edits in multiple samples, allowing to rank sets of gRNAs by activity. This method was also applied to transfected protoplasts (data not shown), making rapid *in-planta* screening of the efficiency of gRNAs possible prior to regeneration of large collections of plant materials. Some samples contain an edit in only a single gene (*e.g.* samples S11, S14, S17), while other samples have edits in multiple genes (*e.g.* up to 11 genes edited in sample S18). HiPlex sequencing of the targeted loci and SMAP analysis thus enables screening of multiple samples and loci in parallel. Importantly, reporting the exact detected nucleotide sequences at each locus allows to interpret the effect of single nucleotide substitutions but also of insertions or deletions (range -21 to +60 nucleotides), onto the encoded protein in the broader genic context.

| | | cultivar | Bintje | | | | | | | | | | | | | | | Spunta | | | |
|---|---|---|---|---|---|---|---|---|---|---|---|---|---|---|---|---|---|---|---|---|---|
| | | # calli/sample | 5 | | | | | | | | | | 1 | | | | | 1 | | | |
| | | pool of gRNA vectors | control | pool 2 | | pool 3 | | | | | pool 4 | | pool 2 | | | pool 1+2 | | control | pool 1+2 | | |
| | | sample | S01 | S02 | S03 | S04 | S05 | S06 | S07 | S08 | S09 | S10 | S11 | S12 | S13 | S14 | S15 | S16 | S17 | S18 | S19 |
| | | # gRNAs/transfection | 0 | 11 | 11 | 11 | 11 | 11 | 11 | 11 | 11 | 11 | 11 | 11 | 11 | 22 | 22 | 0 | 22 | 22 | 22 |
| | | genes edited | 0 | 2 | 2 | 4 | 3 | 2 | 4 | 2 | 5 | 3 | 1 | 4 | 2 | 1 | 7 | 0 | 1 | 11 | 3 |
| Locus | Haplotypes | wt/edit | | | | | | | | | | | | | | | | | | | |
| CPR5_2 | AACTTAAAGAAAGGCAGCTGGCTCTCAGTTCAGATGCGAACCTTTTGGAGAGGGTCAAACTATC.ATTTGGTTTCTCTAAG | wt | 100 | 100 | 100 | 100 | 100 | 100 | 100 | 100 | 100 | 100 | 100 | 100 | | 100 | 100 | 100 | 82 | 65 | 44 |
| CPR5_2 | AACTTAAAGAAAGGCAGCTGGCTCTCAGTTCAGATGCGAACCTTTTGGAGAGGGTCAAA[+60]ATTTGGTTTCTCTAAG | +60 | | | | | | | | | | | | | | | | | | 16 | |
| CPR5_2 | AACTTAAAGAAAGGCAGCTGGCTCTCAGTTCAGATGCGAACCTTTTGGAGAGGGTCAAACTATCAATTTGGTTTCTCTAAG | +1 | | | | | | | | | | | | | | | | | | 6 | |
| CPR5_2 | AACTTAAAGAAAGGCAGCTGGCTCTCAGTTCAGATGCGAACCTTTTGGAGAGGGTCAAACTAT-.ATTTGGTTTCTCTAAG | -1 | | | | | | | | | | | | | | | | | | 13 | |
| CPR5_2 | AACTTAAAGAAAGGCAGCTGGCTCTCAGTTCAGATGCGAACCTTTTGGAGAGGGTCAAAC----.ATTTGGTTTCTCTAAG | -4 |  |  |  |  |  |  |  |  |  |  |  |  |  |  |  |  |  |  | 36 |
| CPR5_2 | AACTTAAAGAAAGGCAGCTGGCTCTCAGTTCAGATGCGAACCTTTTGGAGAGGGTCAAACT---.ATTTGGTTTCTCTAAG | -3 | | | | | | | | | | | | | | | | | 18 | | 20 |
| BON3_3 | AATCCAAATAATCAATACACTTTCAGAAATGCAAGTCACCCTACTGAACTCCGGCTACAGGCATTCATCCCGGACTATATGGGCAAG | wt | 23 | 23 | 21 | 22 | 23 | 23 | 25 | 21 | 23 | 25 | 23 | 24 | 19 | 23 | 25 | 25 | 13 | 12 | 24 |
| BON3_3 | AATCCAAATAATCAATACACTTACAGAAATGCAAGTCACCCTACTGAACTCCGGCTACAGGCATTCATCCCGGACTATATGGGCAAG | wt | 26 | 28 | 32 | 29 | 27 | 30 | 30 | 27 | 29 | 26 | 28 | 29 | 37 | 34 | 29 | 49 | 59 | 14 | 50 |
| BON3_3 | AATCCAAATAACCAATACACTTTCAGAAATGCAAGTCACCCTACTGAACTCCGGCTACAGGCATTCATCCCGGACTATATGGGCAAG | wt | 29 | 25 | 26 | 23 | 26 | 25 | 24 | 26 | 23 | 26 | 26 | 24 | 21 | 23 | 20 | | | | |
| BON3_3 | AATCCAAATAACCAATACACTTTCAGAAATGCAAGTCACCCTACTGAACTCCGGCTACAGTCATTCATCCCGGACTATATGGGCAAG | wt | 21 | 25 | 21 | 26 | 24 | 22 | 21 | 26 | 25 | 22 | 23 | 23 | 23 | 20 | 26 | 26 | 27 | 24 | 26 |
| BON3_3 | AATCCAAATAATCAATACACTTACAGAAATGCAAGTCACCCTACT-------GGCTACAGGCATTCATCCCGGACTATATGGGCAAG | -7 |  |  |  |  |  |  |  |  |  |  |  |  |  |  |  |  |  | 29 |  |
| BON3_3 | AATCCAAATAATCAATACACTTTCAGAAATGCAAGTCACCCTACT-----CCGGCTACAGGCATTCATCCCGGACTATATGGGCAAG | -5 |  |  |  |  |  |  |  |  |  |  |  |  |  |  |  |  |  | 20 |  |
| PMR4_1 | TTCTGCAGTCGTTGCTTGATGCAGGAATGCAATATCGTATCATCT.CTAGGGAGACCCCGTGGCATGGGGTGAGAATGGTGTTGAAGAG | wt | 100 | 100 | 100 | 100 | 100 | 100 | 100 | 100 | 100 | 100 | 100 | 100 | 100 | 76 | 100 | 100 | 100 | 46 | 100 |
| PMR4_1 | TTCTGCAGTCGTTGCTTGATGCAGGAATGCAATATCGTATCATCTTCTAGGGAGACCCCGTGGCATGGGGTGAGAATGGTGTTGAAGAG | +1 | | | | | | | | | | | | | | | | | | 16 | |
| PMR4_1 | TTCTGCAGTCGTTGCTTGATGCAGGAATGCAATATCGTATC---T.CTAGGGAGACCCCGTGGCATGGGGTGAGAATGGTGTTGAAGAG | -3 | | | | | | | | | | | | | | | | | | 16 | |
| PMR4_1 | TTCTGCAGTCGTTGCTTGATGCAGGAATGCAATATCGTATC----.CTAGGGAGACCCCGTGGCATGGGGTGAGAATGGTGTTGAAGAG | -4 |  |  |  |  |  |  |  |  |  |  |  |  |  |  |  |  |  | 14 |  |
| PMR4_1 | TTCTGCAGTCGTTGCTTGATGCAGGAATGC---------------.--------ACCCCGTGGCATGGGGTGAGAATGGTGTTGAAGAG | -23 |  |  |  |  |  |  |  |  |  |  |  |  |  |  |  |  |  | 8 |  |
| PMR4_1 | TTCTGCAGTCGTTGCTTGATGCAGGAATGCAATATCGTATCATCTACTAGGGAGACCCCGTGGCATGGGGTGAGAATGGTGTTGAAGAG | +1 | | | | | | | | | | | | | 24 | | | | | | |
| PSY1R_3 | AAAACTGATGGAGCATCCCTACTTGATTGGCCAACTCGACTGAAGATTGCAC.AGGGGGCAAGTTGTGGGCTGGCTTACATGCATCAGATATGTGA | wt | 72 | 69 | 73 | 76 | 73 | 75 | 81 | 74 | 70 | 75 | 75 | 72 | 69 | 75 | 56 | 14 | 27 | | 25 |
| PSY1R_3 | AAAACTGATGGAGCATTCCTACTTGATTGGCCAACTCGACTGAAGATTGCAC.AGGGGGCAAGTTGTGGGCTGGCTTACATGCATCAGATATGTGA | wt | 28 | 31 | 27 | 24 | 27 | 25 | 19 | 26 | 30 | 25 | 25 | 28 | 31 | 25 | 31 | 86 | 73 | 50 | 75 |
| PSY1R_3 | AAAACTGATGGAGCATTCCTACTTGATTGGCCAACTCGACTGAAGATTGC--.---GGGCAAGTTGTGGGCTGGCTTACATGCATCAGATATGTGA | -5 | | | | | | | | | | | | | | | | | | 23 | |
| PSY1R_3 | AAAACTGATGGAGCATTCCTACTTGATTGGCCAACTCGACTGAAGATTGCACCAGGGGGCAAGTTGTGGGCTGGCTTACATGCATCAGATATGTGA | +1 | | | | | | | | | | | | | | | | | | 27 | |
| PSY1R_3 | AAAACTGATGGAGCATCCCTACTTGATTGGCCAACTCGACTGAAGATTGC--.---GGGCAAGTTGTGGGCTGGCTTACATGCATCAGATATGTGA | -5 | | | | | | | | | | | | | | | 13 | | | | |
| LACS2_2 | AAGTTGTGGAGGATGTCTCACATCCATAGCCAATGTACACTCCATGACAGGAAC.TGTTGGTGTTCCAATGACTACAATTGAGGCAAGACTT | wt | 100 | 93 | | | 100 | 100 | | 100 | 100 | 100 | 100 | 100 | 100 | 100 | 45 | 100 | 100 | 2 | |
| LACS2_2 | AAGTTGTGGAGGATGTCTCACATCCATAGCCAATGTACACTCCATGACAGGAAC.TGTTTGTGTTCCAATGACTACAATTGAGGCAAGACTT | snp | | 8 | | | | | | | | | | | | | | | | | |
| LACS2_2 | AAGTTGTGGAGGATGTCTCACATCCATAGCCAATGTACACTCCATGACAGG--C.TGTTGGTGTTCCAATGACTACAATTGAGGCAAGACTT | -2 | | | | | | | | | | | | | | | | | | 40 | |
| LACS2_2 | AAGTTGTGGAGGATGTCTCACATCCATAGCCAATGTACACTCCATGACAGGAA-.TGTTGGTGTTCCAATGACTACAATTGAGGCAAGACTT | -1 | | | | | | | | | | | | | | | | | | 20 | |
| LACS2_2 | AAGTTGTGGAGGATGTCTCACATCCATAGCCAATGTACACTCCATGACA-----.--TTGGTGTTCCAATGACTACAATTGAGGCAAGACTT | -7 |  |  |  |  |  |  |  |  |  |  |  |  |  |  |  |  |  | 38 |  |
| LACS2_2 | AAGTTGTGGAGGATGTCTCACATCCATAGCCAATGTACACTCCATGACAGGAACATGTTGGTGTTCCAATGACTACAATTGAGGCAAGACTT | +1 | | | | | | | | | | | | | | | 13 | | | | |
| LACS2_2 | AAGTTGTGGAGGATGTCTCACATCCATAGCCAATGTACACTCCATGACAGG---.TGTTGGTGTTCCAATGACTACAATTGAGGCAAGACTT | -4 | | | | | | | | | | | | | | | 25 | | | | |
| LACS2_2 | AAGTTGTGGAGGATGTCTCACATCCATAGCCAATGTACACTCCATGACAG----.TGTTGGTGTTCCAATGACTACAATTGAGGCAAGACTT | -5 |  |  |  |  |  |  |  |  |  |  |  |  |  |  | 16 |  |  |  |  |
| LCR_1 | TGACACGTGGTTATTCCAGCGCAAGACTGCTGCGCTGGAATTCACCACGCGGACC.TTGAGGCAGGCTATGGGTCGATGGGTGAACCGAGC | wt | 54 | 51 | 53 | 49 | 54 | 52 | 43 | 49 | 53 | 51 | 51 | 57 | 55 | 55 | 59 | 56 | 59 | 38 | 56 |
| LCR_1 | TGACACGTGGTTATTCCAGCGCAAAACTGCTGCGCTGGAATTCACCACGCGGACC.TTGAGGCAGGCTATGGGTCGATGGGTGAACCGAGC | wt | 46 | 49 | 47 | 51 | 46 | 48 | 57 | 51 | 47 | 49 | 49 | 43 | 45 | 45 | 35 | 44 | 41 | 32 | 44 |
| LCR_1 | TGACACGTGGTTATTCCAGCGCAAAACTGCTGCGCTGGAATTCACCACGCGGACCTTTGAGGCAGGCTATGGGTCGATGGGTGAACCGAGC | +1 | | | | | | | | | | | | | | | 6 | | | | |
| LCR_1 | TGACACGTGGTTATTCCAGCGCAAAACTGCTGCGCTGGAATTCACCACGCGGACC.-TGAGGCAGGCTATGGGTCGATGGGTGAACCGAGC | -1 | | | | | | | | | | | | | | | | | | 13 | |
| LCR_1 | TGACACGTGGTTATTCCAGCGCAAGACTGCTGCGCTGGAATTCACCACGCGGACCCTTGAGGCAGGCTATGGGTCGATGGGTGAACCGAGC | +1 | | | | | | | | | | | | | | | | | | 17 | |
| CPR1_1 | AAGGTGTTTAGGAAGTTACCCTTTTGTCAGTTTAACCCTCCGAGGAAACCCCCGGGTGGACGGGGTCCCGGGTTAGCTCAGATTCGTG | wt | 96 | 94 | 92 | 94 | 95 | 91 | 91 | 95 | 96 | 94 | 95 | 97 | 96 | 97 | 64 | 97 | 97 | | |
| CPR1_1 | AAGGTATTTAGGAAGTTACCCTTTTGTCAGTTTAACCCTCCGAGGAAACCCCCGGGTGGACGGGGTCCCGGGGTTGGCTCAGATTCGTG | snp | 4 | 6 | 8 | 6 | 5 | 9 | 9 | 5 | 4 | 6 | 5 | 3 | 4 | 3 | 5 | 3 | 3 | | |
| CPR1_1 | AAGGTGTTTAGGAAGTTACCCTTTTGTCAGTTTAACCCTCCGAGGAA--CCCCGGGTGGACGGGGTCCCGGGTTAGCTCAGATTCGTG | -2 | | | | | | | | | | | | | | | 26 | | | 50 | 100 |
| CPR1_1 | AAGGTGTTTAGGAAGTTACCCTTTTGTCAGTTTAACCCTCCGAGGA---CCCCGGGTGGACGGGGTCCCGGGTTAGCTCAGATTCGTG | -3 | | | | | | | | | | | | | | | 4 | | | 50 | |
| CPR1_2 | AGGTTCAGGATTGTCCTTACTGGTTGCTTAAAGAGGATAATGGGGCTTG.TGCTGGAGGTGCATTGCATTGGATTGTAACGAAGGAACCG | wt | 71 | 75 | 68 | 81 | 77 | 74 | 80 | 79 | 77 | 78 | 74 | 78 | 77 | 75 | 54 | 72 | 74 | 61 | 100 |
| CPR1_2 | AGGTTCAGGATTGTCCTTACTGGTTGCTTAAGGAGGATAATGGGGCTTG.TGCTGGAGGTGCATTGCATTGGATTGTAACGAAGGAACCG | wt | 29 | 25 | 32 | 19 | 23 | 26 | 20 | 21 | 23 | 22 | 26 | 15 | 23 | 25 | 28 | 28 | 26 | 21 | |
| CPR1_2 | AGGTTCAGGATTGTCCTTACTGGTTGCTTAAGGAGGATAATGGGGCTTGTTGCTGGAGGTGCATTGCATTGGATTGTAACGAAGGAACCG | +1 | | | | | | | | | | | | | | | | | | 19 | |
| CPR1_2 | AGGTTCAGGATTGTCCTTACTGGTTGCTTAAAGAGGATAATGGGGCTTGTTGCTGGAGGTGCATTGCATTGGATTGTAACGAAGGAACCG | +1 | | | | | | | | | | | | | | | 18 | | | | |
| CPR1_2 | AGGTTCAGGATTGTCCTTACTGGTTGCTTAAGGAGGATAATGGGGCTTG.-GCTGGAGGTGCATTGCATTGGATTGTAACGAAGGAACCG | -1 | | | | | | | | | | | 8 | | | | | | | | |
| PUB24_2 | TAAGATCATCGAATCAGGAGCCATATTCGAGCTCATAGAGCTCGA.ATTAGGCAACACTGAAAAGAGGGTTTCAGAGTTGATACTTTGTGTTCTAG | wt | 23 | 24 | 24 | 28 | 26 | 25 | 22 | 25 | 25 | 25 | 20 | 21 | 26 | 27 | 30 | 26 | 27 | | 7 |
| PUB24_2 | TAAGATCATCGAATCAGGAGCTATATTCGAGCTCATAGAGCTCGA.ATTAGGCAACACTGAAAAGAGGGTTTCAGAGTTGGTACTTTGTGTTCTAG | wt | 27 | 24 | 28 | 25 | 26 | 28 | 27 | 27 | 25 | 25 | 27 | 24 | 22 | 26 | 30 | 24 | 24 | 22 | |
| PUB24_2 | TAAGATCATCGAATCAGGAGCCATATTCGAGCTCATAGAGCTCGA.ATTAGGCAACACTGAAAAGAGGGTTTCAGAGTTGGTACTTTGTGTTCTAG | wt | 50 | 52 | 48 | 47 | 48 | 48 | 51 | 48 | 50 | 51 | 53 | 50 | 51 | 48 | 30 | 50 | 50 | 56 | 49 |
| PUB24_2 | TAAGATCATCGAATCAGGAGCCATATTCGAGCTCATAGAGCTCG-.ATTAGGCAACACTGAAAAGAGGGTTTCAGAGTTGATACTTTGTGTTCTAG | -1 | | | | | | | | | | | | 5 | | | | | | 22 | |
| PUB24_2 | TAAGATCATCGAATCAGGAGCCATATTCGAGCTCATAGAGCTCG-.ATTAGGCAACACTGAAAAGAGGGTTTCAGAGTTGGTACTTTGTGTTCTAG | -1 | | | | | | | | | | | | | | | | | | | 43 |
| PUB24_2 | TAAGATCATCGAATCAGGAGCCATATTCGAGCTCATAGAGCTCGAGATTAGGCAACACTGAAAAGAGGGTTTCAGAGTTGGTACTTTGTGTTCTAG | +1 | | | | | | | | | | | | | | | 11 | | | | |
| MLO1_3 | TTCTAATCCAGTGACACCAATGTCAAGTAGACCAGGGACACCTTCTCA-TCATGGCATGTCACCCGTGCATTTATTGCGCGGACACTATA | wt | 100 | 89 | 88 | 100 | 100 | 100 | 100 | 100 | 100 | 100 | 100 | 89 | 67 | 100 | 100 | 100 | 100 | 85 | 100 |
| MLO1_3 | TTCTAATCCAGTGACACCAATGTCAAGTAGACCAGGGACACCTTCTCAATCATGGCATGTCACCCGTGCATTTATTGCGCGGACACTATA | +1 | | | | | | | | | | | | | 33 | | | | | | |
| MLO1_3 | TTCTAATCCAGTGACACCAATGTCAAGTAGACCAGGGACACCTTCTCACTCATGGCATGTCACCCGTGCATTTATTGCGCGGACACTATA | +1 | | 11 | | | | | | | | | | | | | | | | | |
| MLO1_3 | TTCTAATCCAGTGACACCAATGTCAAGTAGACCAGGGACACCTTC---.TCATGGCATGTCACCCGTGCATTTATTGCGCGGACACTATA | -3 | | | 12 | | | | | | | | | 11 | | | | | | | |
| MLO1_3 | TTCTAATCCAGTGACACCAATGTCAAGTAGACCAGGGA----------.-CATGGCATGTCACCCGTGCATTTATTGCGCGGACACTATA | -11 |  |  |  |  |  |  |  |  |  |  |  |  |  |  |  |  |  |  |  |
| MLO1_3 | TTCTAATCCAGTGACACCAATGTCAAGTAGACCAGGGACACCTT----.--ATGGCATGTCACCCGTGCATTTATTGCGCGGACACTATA | -6 |  |  |  |  |  |  |  |  |  |  |  |  |  |  |  |  |  | 15 |  |
| EXLA2_2 | CTTGACCCGAAACCACGGAGCAATTTGGGACACAAGCAGAGTGCCA.AGTGGGGCATTGCAATTTAGGTTTGTGGTGACAGCAGGGT | wt | 100 | 45 | 59 | 100 | 100 | 100 | 100 | 100 | 100 | 100 | 53 | 55 | | 100 | 29 | 100 | 100 | | |
| EXLA2_2 | CTTGACCCGAAACCACGGAGCAATTTGGGACACAAGCAGAGTGCC-.AGTGGGGCATTGCAATTTAGGTTTGTGGTGACAGCAGGGT | -1 | | 12 | | | | | | | | | 19 | 20 | 46 | | 71 | | | | |
| EXLA2_2 | CTTGACCCGAAACCACGGAGCAATTTGGGACACAAGCAGAGTGCCAAAGTGGGGCATTGCAATTTAGGTTTGTGGTGACAGCAGGGT | +1 | | 26 | 5 | | | | | | | | | 14 | | | | | | | |
| EXLA2_2 | CTTGACCCGAAACCACGGAGCAATTTGGGACACAAGCAGAGT----.AGTGGGGCATTGCAATTTAGGTTTGTGGTGACAGCAGGGT | -4 |  | 4 |  |  |  |  |  |  |  |  | 12 |  | 20 |  |  |  |  | 22 |  |
| EXLA2_2 | CTTGACCCGAAACCACGGAGCAATTTGGGACACAAGCAG-------.AGTGGGGCATTGCAATTTAGGTTTGTGGTGACAGCAGGGT | -7 |  | 7 |  |  |  |  |  |  |  |  |  |  |  |  |  |  |  |  |  |
| EXLA2_2 | CTTGACCCGAAACCACGGAGCAATTTGGGACACAAGCAGAGTGCCA.------GCATTGCAATTTAGGTTTGTGGTGACAGCAGGGT | -6 |  | 6 |  |  |  |  |  |  |  |  |  |  |  |  |  |  |  |  |  |
| EXLA2_2 | CTTGACCCGAAACCACGGAGCAATTTGGGACACAAGCAGAGTGCC-.----------TGCAATTTAGGTTTGTGGTGACAGCAGGGT | -11 |  |  | 15 |  |  |  |  |  |  |  |  |  |  |  |  |  |  |  |  |
| EXLA2_2 | CTTGACCCGAAACCACGGAGCAATTTGGGACACAAGCAGAGTGC--.AGTGGGGCATTGCAATTTAGGTTTGTGGTGACAGCAGGGT | -2 | | | 21 | | | | | | | | | | | | | | | 42 | |
| EXLA2_2 | CTTGACCCGAAACCACGGAGCAATTTGGGACACAAGCAGAGTGCCATAGTGGGGCATTGCAATTTAGGTTTGTGGTGACAGCAGGGT | +1 | | | | | | | | | | | | | | | | | | 35 | |
| EXLA2_2 | CTTGACCCGAAACCACGGAGCAATTTGGGACACAAGCAGA----CA.AGTGGGGCATTGCAATTTAGGTTTGTGGTGACAGCAGGGT | -4 |  |  |  |  |  |  |  |  |  |  | 16 |  |  |  |  |  |  |  |  |
| EXLA2_2 | CTTGACCCGAAACCACGGAGCAATTTGGGACACAAGCAGAG-----.AGTGGGGCATTGCAATTTAGGTTTGTGGTGACAGCAGGGT | -5 |  |  |  |  |  |  |  |  |  |  |  | 6 |  |  |  |  |  |  |  |
| EXLA2_2 | CTTGACCCGAAACCACGGAGCAATTTGGGACACAAGCAGAGTGCCA[+21]GTGGGGCATTGCAATTTAGGTTTGTGGTGACAGCAGGGT | +21 | | | | | | | | | | | | 5 | | | | | | | |
| EXLA2_2 | CTTGACCCGAAACCACGGAGCAATTTGGGACACAAG----------.AGTGGGGCATTGCAATTTAGGTTTGTGGTGACAGCAGGGT | -10 |  |  |  |  |  |  |  |  |  |  |  |  | 34 |  |  |  |  |  |  |
| LSD1_1 | TGTCGTACATTACTTATGCATCCACGTGGAGCGAACAGTGTGAGATGCTCCTGCTGTC.ATACGGTGAACCTTGTCCCAGGTACAGTTTATAACT | wt | 100 | 100 | 100 | 85 | 95 | 82 | 84 | 75 | 100 | 100 | 100 | 100 | 100 | 100 | 100 | 100 | 100 | 100 | 100 |
| LSD1_1 | TGTCGTACATTACTTATGCATCCACGTGGAGCGAACAGTGTGAGATGC----------.-TACGGTGAACCTTGTCCCAGGTACAGTTTATAACT | -11 |  |  |  | 7 |  |  |  |  |  |  |  |  |  |  |  |  |  |  |  |
| LSD1_1 | TGTCGTACATTACTTATGCATCCACGTGGAGCGAACAGTGTGAGATGCTCCTGCTGTCAATACGGTGAACCTTGTCCCAGGTACAGTTTATAACT | +1 | | | | 8 | 5 | 11 | 16 | 25 | | | | | | | | | | | |
| LSD1_1 | TGTCGTACATTACTTATGCATCCACGTGGAGCGAACAGTGTGAGATGCTCCTGCT---.ATACGGTGAACCTTGTCCCAGGTACAGTTTATAACT | -3 | | | | | | 7 | | | | | | | | | | | | | |
| EDR2_3 | ACTGGTTCAAGGATACTAAACGAATGGACCATGTC.GCTAGGCGTCCTGGTTGTGCAGCACAGGTAAGTTTAAAGGGAATGTCAC | wt | 51 | 47 | 47 | 50 | 37 | 46 | 50 | 48 | 49 | 47 | 48 | 49 | 46 | 47 | 56 | 53 | 50 | 50 | 52 |
| EDR2_3 | ATTGGTTCAAGGATACTAAACGAATGGACCATGTC.GCTAGGCATCCTGGCTGTGCAGCACAGGTAAGTTTAAAGGGAATGTCAA | wt | 30 | 28 | 31 | 23 | 22 | 25 | 26 | 27 | 28 | 28 | 30 | 28 | 30 | 30 | 25 | | | | |
| EDR2_3 | ACTGGTTCAAGGATACTAAACGAATGGACCATGTC.GCTAGGCGTCCTGGTTGTGCAGCACAGGTAAGTTTAAAGGGAATGCCAA | wt | 19 | 24 | 22 | 22 | 20 | 24 | 15 | 25 | 23 | 25 | 22 | 23 | 25 | 23 | 19 | 47 | 50 | 50 | 48 |
| EDR2_3 | ATTGGTTCAAGGATACTAAACGAATGGACCATGTCCGCTAGGCATCCTGGCTGTGCAGCACAGGTAAGTTTAAAGGGAATGTCAA | +1 | | | | 6 | | | | | | | | | | | | | | | |
| EDR2_3 | ACTGGTTCAAGGATACTAAACGAATGGACCATGTCAGCTAGGCGTCCTGGTTGTGCAGCACAGGTAAGTTTAAAGGGAATGTCAC | +1 | | | | | 10 | | | | | | | | | | | | | | |
| EDR2_3 | ATTGGTTCAAGGATACTAAACGAATGGACCATGTTCGCTAGGCATCCTGGCTGTGCAGCACAGGTAAGTTTAAAGGGAATGTCAA | +1 | | | | | 6 | | | | | | | | | | | | | | |
| EDR2_3 | ACTGGTTCAAGGATACTAAACGAATGGACCATGTCTGCTAGGCGTCCTGGTTGTGCAGCACAGGTAAGTTTAAAGGGAATGCCAA | +1 | | | | | 6 | | 3 | | | | | | | | | | | | |
| EDR2_3 | ATTGGTTCAAGGATACTAAACGAATGGACCATGTCAGCTAGGCATCCTGGCTGTGCAGCACAGGTAAGTTTAAAGGGAATGTCAA | +1 | | | | | | 5 | 6 | | | | | | | | | | | | |
| FER_1 | CCGTCTGTATTTCTATCCCTCGTCTTACAACAAGCTAAATGCTACTAATGCCATCTTCA.ATGTGGCCTCCGGACAATATACCTTACTTAGAAA | wt | 77 | 70 | 77 | 72 | 71 | 75 | 77 | 74 | 74 | 71 | 76 | 75 | 74 | 73 | 71 | 100 | 100 | 100 | 100 |
| FER_1 | CCGTCTGTATTTCTATCCCTCGTCTTACAACAAGCTAAATGCTACTAATGCCATCTTCA.ATGTGGCCTCCGGACGATATACCTTACTTAGAAA | wt | 23 | 30 | 23 | 21 | 29 | 25 | 23 | 26 | 26 | 29 | 24 | 25 | 26 | 27 | 29 | | | | |
| FER_1 | CCGTCTGTATTTCTATCCCTCGTCTTACAACAAGCTAAATGCTACTAATGCCATCTTCACATGTGGCCTCCGGACAATATACCTTACTTAGAAA | +1 | | | | 7 | | | | | | | | | | | | | | | |
| MYB3R4_1 | TAAGCTAATCGGATTAGTTAACATACATGGCCCCAAAAAATGGTCCACCATTGCACAAGAGTTAGCAGGACGTATTGGAAAACAATGT | wt | 44 | 51 | 52 | 43 | 48 | 50 | 52 | 48 | 48 | 55 | 47 | 49 | 50 | 50 | 55 | 100 | 100 | 100 | 100 |
| MYB3R4_1 | TAAGCTAATTGGATTAGTTAACATACATGGCCCCAAAAAATGGTCCACCATTGCACAAGAGTTAGCAGGACGTATTGGAAAACAATGT | wt | 56 | 49 | 48 | 52 | 53 | 50 | 48 | 52 | 52 | 45 | 53 | 51 | 50 | 50 | 45 | | | | |
| MYB3R4_1 | TAAGCTAATCGGATTAGTTAACATACATGGCC-------------ACCATTGCACAAGAGTTAGCAGGACGTATTGGAAAACAATGT | -13 |  |  |  | 5 |  |  |  |  |  |  |  |  |  |  |  |  |  |  |  |
| PSKR1_1 | TAGTCTTGTTTCGCTTGATCTGGCTACGAACGGGTTCCGTGGGTTAGTTCCTGAATATCTTCCTAACTGTCAAAGGTTGCAAACTATCAATC | wt | 21 | 25 | | | 32 | 24 | 13 | 28 | 38 | 30 | 32 | 30 | 30 | 38 | 36 | | | | |
| PSKR1_1 | TAGTCTTGTTTCGCTTGATCTGGCTACGAACGGGTTCCGTGGGTTAGTTCCTGAATATCTTCCTGACTGTCGAAGGTTGCAAACTATCAATC | wt | 34 | 32 | | | 20 | 33 | 31 | 29 | 24 | 24 | 37 | 28 | 38 | 26 | 21 | 47 | 52 | 38 | 69 |
| PSKR1_1 | TAGTCTTGTTTCGCTTGATCTGGCTACGAACGGGTTCCATGGGTTAGTTCCTGAATATCTTCCTAACTGTCAAAGGTTGCAAACTATCAATC | wt (snp@gRNA) | 46 | 43 | | | 40 | 44 | 49 | 43 | 38 | 46 | 31 | 42 | 32 | 36 | 43 | 10 | 14 | 21 | 31 |
| PSKR1_1 | TAGTCTTGTTTCGCTTGATCTGGCTACAAACGGGTTCCGTGGGTTAGTTCCTGATTATCTTCCTAACTGTCAAAGGTTGCAAACTATCAATC | wt (snp@gRNA) | | | | | | | | | | | | | | | | 43 | 34 | 41 | |
| PSKR1_1 | TAGTCTTGTTTCGCTTGATCTGGCTACGAACGGGTT-CGTGGGTTAGTTCCTGAATATCTTCCTGACTGTCGAAGGTTGCAAACTATCAATC | -1 | | | | | 8 | | | | | | | | | | | | | | |
| PSKR1_1 | TAGTCTTGTTTCGCTTGATCTGGCTACGAACGGGTT-CGTGGGTTAGTTCCTGAATATCTTCCTAACTGTCAAAGGTTGCAAACTATCAATC | -1 | | | | | | | 8 | | | | | | | | | | | | |
| BON1_1 | TCATGCTCCACGACCAAGGCTTGACCGGAGATACTCGAGGATTGCAGACAATTATAATTCCCTGGATGAGGTATGGATTA | wt | 100 | 100 | 100 | 100 | 100 | 100 | 94 | 100 | 100 | 100 | 100 | 100 | 100 | 100 | 100 | 48 | 50 | 52 | 49 |
| BON1_1 | TCATGCTCCACGACCAAGGCTTGACCGGAGATACTCGAGGATTGCAGACAATTATAATTCCCTGGATGAGGTATGGACTA | wt | | | | | | | | | | | | | | | | 26 | 24 | 23 | 23 |
| BON1_1 | TCATGCTCCACGACCAAGGCTTGACCGGAGATACTCGAGGATTGTAGACAATTATAATTCCCTGGATGAGGTATGGATTA | wt | | | | | | | | | | | | | | | | 25 | 26 | 25 | 28 |
| BON1_1 | TCATGCTCCACGACCAAGGCTTGACCGGA----CTCGAGGATTGCAGACAATTATAATTCCCTGGATGAGGTATGGATTA | -4 |  |  |  |  |  |  | 6 |  |  |  |  |  |  |  |  |  |  |  |  |
| CPR22_1 | GAATTTTTAGAATTTATCCATTGTACAAAGAAGTAACAAGGACTTCAGGCTTATTTACTGAAAGTGCATGGGGTGGAGCTGTTTTCAA | wt | 100 | 100 | 100 | 100 | 100 | 100 | 100 | 94 | 100 | 100 | 97 | 100 | 100 | 100 | 100 | 69 | 74 | 72 | 75 |
| CPR22_1 | GAATTTTTAGAATTTATCCATTGTGCAAAGAAGTAACAAGGACTTCAGGCTTATTTACTGAAAGTGCATGGGGTGGAGCTGTTTTCAA | wt | | | | | | | | | | | 3 | | | | | 31 | 26 | 28 | 25 |
| CPR22_1 | GAATTTTTAGAATTTATCCATTGTACAAAGAAGTAACAAG--CTTCAGGCTTATTTACTGAAAGTGCATGGGGTGGAGCTGTTTTCAA | -2 | | | | | | | | 6 | | | | | | | | | | | |
| BDG_1 | TACAAAATCAAAGTATCGATTATTCGCGGTGGATCT.ACTAGGATTTGGAAGAAGTCCGAAGCCAAATGAGTCACTTTA | wt | 100 | 100 | 100 | 100 | 100 | 100 | 100 | 100 | 77 | 90 | 100 | 100 | 100 | 100 | 100 | 100 | 100 | 100 | 100 |
| BDG_1 | TACAAAATCAAAGTATCGATTATTCGCGGTGGATCTTACTAGGATTTGGAAGAAGTCCGAAGCCAAATGAGTCACTTTA | +1 | | | | | | | | | 15 | | | | | | | | | | |
| BDG_1 | TACAAAATCAAAGTATCGATTATTCGCGGTG----T.ACTAGGATTTGGAAGAAGTCCGAAGCCAAATGAGTCACTTTA | -4 |  |  |  |  |  |  |  |  | 8 |  |  |  |  |  |  |  |  |  |  |
| BDG_1 | TACAAAATCAAAGTATCGATTATTCGCGGTGGAT--.ACTAGGATTTGGAAGAAGTCCGAAGCCAAATGAGTCACTTTA | -2 | | | | | | | | | | 10 | | | | | | | | | |
| BON3_1 | CAGCTTCTATCCCGATGAGAAATTTTGTAATGGATTCGAGGAAGTACTGGGTAGATATAGAGAATTAGTTCCCCAATTACG | wt | 34 | 38 | | | 42 | 47 | | | 43 | 55 | 42 | 57 | 45 | 57 | 54 | 34 | 25 | 47 | 20 |
| BON3_1 | CAGCTTCTATCCCGATGAGAAATTTTGTAATGGATTCGAGGAAGTACTGGGTAGATATAGAGAATTAGTTCCTCAATTACG | wt | 28 | 26 | | | 30 | 20 | | | 29 | 36 | 34 | 22 | 31 | 20 | 32 | 42 | 52 | 45 | 51 |
| BON3_1 | CAGCTTCTATCCCGATGAGAAATTTTGTAATGGATTCGAGGAAGTACTGGGTAGATATAGAGAATTAGTTCCCCAGTTACG | wt | 38 | 35 | | | 28 | 33 | | | 20 | 9 | 24 | 21 | 24 | 23 | 14 | 24 | 24 | 8 | 30 |
| BON3_1 | CAGCTTCTATCCCGATGAGAAATTTTGTAATGGATTCGAGGAAG[+58]TACTGGGTAGATATAGAGAATTAGTTCCCCAGTTACG | +58 | | | | | | | | | 9 | | | | | | | | | | |
| CPR5_1 | ATGTCATGAATCAATAGGTCGGCAGCTTACCTTCTATGATAGGCAA.GTAAGGCAGCAATTGGCTTCTGCTTCTTCAAGCATGATT | wt | 75 | 76 | 77 | 78 | 76 | 78 | 76 | 77 | 74 | 68 | 73 | 76 | 74 | 73 | 82 | 30 | 31 | 28 | 24 |
| CPR5_1 | ATGTCGTGAATCAATAGGTCGGCAGCTTACCTTCTATGATAGGCAA.GTAAGGCAGCAATTGGCTTCTGCTTCTTCAAGCATGATT | wt | 25 | 24 | 23 | 22 | 24 | 22 | 24 | 23 | 26 | 24 | 27 | 24 | 26 | 27 | 18 | 70 | 69 | 72 | 76 |
| CPR5_1 | ATGTCATGAATCAATAGGTCGGCAGCTTACCTTCTATGATAGGCAAAGTAAGGCAGCAATTGGCTTCTGCTTCTTCAAGCATGATT | +1 | | | | | | | | | | 7 | | | | | | | | | |
| DMR1_1 | GTCGTTTGCTCCGGCCACTGTTGCTAATCTAGG.TCCGGGTTTTGATTT-CCTCGGATGCGCCGTTGATGGAATCGGAGATT | wt | 79 | 75 | 73 | 73 | 73 | 74 | 74 | 76 | 71 | 77 | 76 | 77 | 74 | 74 | 79 | 36 | 29 | 25 | 27 |
| DMR1_1 | GTCGTTTGCTCCGGCCACTGTTGCTAATCTAGG.TCCGGGTTTTGATTTTCTT-GGATGCGCCGTTGATGGAATCGGAGATT | wt | 21 | 25 | 27 | 27 | 27 | 26 | 26 | 24 | 23 | 23 | 24 | 23 | 26 | 26 | 21 | 52 | 46 | 48 | 48 |
| DMR1_1 | GTCGTTTGCTCCGGCCACTGTTGCTAATCTAGG.TCCGGGTTTTGATTT-CCTCGGATGCGCTGTTGATGGAATCGGAGATT | wt | | | | | | | | | | | | | | | | 12 | 24 | 27 | 25 |
| DMR1_1 | GTCGTTTGCTCCGGCCACTGTTGCTAATCTAGGTTCCGGGTTTTGATTT-CCTCGGATGCGCCGTTGATGGAATCGGAGATT | +1 | | | | | | | | | 7 | | | | | | | | | | |
| FER_3 | GATTTTGGTTTGTCTAAAACAGGTCCTACATTGGATCACACCCATGTCA.GTACGGTGGTGAAGGGCAGTTTTGGATATCTGGATCCAGAATACT | wt | 100 | 100 | 100 | 100 | 100 | 100 | 100 | 100 | 73 | 100 | 100 | 100 | 100 | 100 | 100 | 86 | 85 | 86 | 85 |
| FER_3 | GATTTTGGGTTGTCTAAAACAGGCCCTACATTGGATCACACCCATGTCA.GTACCGTGGTGAAGGGCAGTTTTGGATATCTGGATCCAGAATACT | wt (snp@PAM) | | | | | | | | | | | | | | | | 14 | 15 | 14 | 15 |
| FER_3 | GATTTTGGTTTGTCTAAAACAGGTCCTACATTGGATCACACCCATGTCAAGTACGGTGGTGAAGGGCAGTTTTGGATATCTGGATCCAGAATACT | +1 | | | | | | | | | 27 | | | | | | | | | | |
| WAT1_3 | GGTATAGCATTTGCAGTACAGATATGGTGCATTGACA.GAGGGGGTCCAGTTTTTGTTGCTGTTTATCAACCTGTTCAGACTCTTGTTGTTGCTC | wt | 78 | 80 | 80 | 81 | 75 | 75 | 84 | 81 | 45 | 53 | 74 | 79 | 79 | 77 | 80 | 100 | 100 | 100 | 100 |
| WAT1_3 | GGTATAGCATTTGCAGTACAGATATGGTGCATTGACA.GAGGGGGTCCAGTCTTTGTTGCTGTTTATCAACCTGTTCAGACTCTTGTTGTTGCTC | wt | 22 | 20 | 20 | 19 | 25 | 25 | 16 | 19 | 15 | 25 | 26 | 21 | 21 | 23 | 20 | | | | |
| WAT1_3 | GGTATAGCATTTGCAGTACAGATATGGTGCATTGACAAGAGGGGGTCCAGTTTTTGTTGCTGTTTATCAACCTGTTCAGACTCTTGTTGTTGCTC | +1 | | | | | | | | | 33 | 15 | | | | | | | | | |
| WAT1_3 | GGTATAGCATTTGCAGTACAGATATGGTGCATTG---.GAGGGGGTCCAGTTTTTGTTGCTGTTTATCAACCTGTTCAGACTCTTGTTGTTGCTC | -3 | | | | | | | | | 7 | | | | | | | | | | |
| WAT1_3 | GGTATAGCATTTGCAGTACAGATATGGTGCA------.GAGGGGGTCCAGTTTTTGTTGCTGTTTATCAACCTGTTCAGACTCTTGTTGTTGCTC | -6 |  |  |  |  |  |  |  |  |  | 7 |  |  |  |  |  |  |  |  |  |
| CPR5_2 : [+60] | TGCAGTGCTGCCATAACCATGAGTGATAACACTGCGGCCAACTTACTTCTGACAACGATC (fragment of backbone gRNA expression vector) | | | | | | | | | | | | | | | | | | | | |
| EXLA2_2 : [+21] | TTCGCCACCTCTGACTTGAGC (fragment of backbone Cas9 expression vector) | | | | | | | | | | | | | | | | | | | | |
| BON3_1 : [+58] | ATCGCTGTTAAAAGGACAATTACAAACAGGAATCGAATGCAACCGGCGCAGGAACACT (fragment of backbone Cas9 expression vector) | | | | | | | | | | | | | | | | | | | | |

**Figure S7 | Multiplex CRISPR/Cas genome editing in potato.** *SMAP haplotype-window* applied to CRISPR-induced mutation screens and allelic variation in two potato cultivars. Column "Locus" shows target genes and amplicon number. Column "Haplotype" lists the four endogenous haplotypes of tetraploid potato and any edited haplotypes per locus. gRNAs target sequences are indicated in orange, with the PAM site marked in bold. Edits and SNPs are marked in red, single nucleotide insertions are aligned with a red '.', while deletions are aligned with a '-'. Column "wt/edit" lists the length difference with the respective wild type (wt) reference haplotype. The genotype call table shows relative frequency per haplotype per locus for each sample (S01-S19). Control samples (transfected with Cas9 only, no gRNA vector), show wild type (non-edited) haplotypes for cultivars Bintje (S01) and Spunta (S16). Samples are either pools of five calli (Bintje), or individual calli (Bintje and Spunta). gRNA expression vectors are pooled for co-transfection with the Cas9 expression vector, in subsets of 11 vectors (pool 2, 3, 4), or 22 vectors (pool 1+2). Per sample, the number of edited genes is listed. Edits are consistently located at -4/-3 bp from the PAM site indicating Cas9 activity. Large insert sequences for CPR5_2 [+58], EXLA2_2 [+21], BON3_1 [+60] are shown below the table, and correspond to Cas9/gRNA vector fragments. The HiPlex assay simultaneously screens 222 amplified fragments across 88 genes, but only loci with edits are shown.

#### Affiliations

[1]Flanders Research Institute for Agriculture, Fisheries and Food, Plant Sciences Unit, Caritasstraat 39, B-9090, Melle, Belgium, [2]Ghent University, Department of Applied Mathematics, Computer Science and Statistics, Krijgslaan 281 S9, B-9000, Ghent, Belgium, [3]INRAE, URP3F, F-86600 Lusignan, France, [4]Teagasc, Crop Science Department, Oak Park, Carlow R93 XE12, Ireland. Correspondence should be addressed to T.R..

#### Methods

Methods, including a detailed user manual explaining the features and optional parameters of the code, statements of data availability and any associated accession codes and references, are available in the **Online manual**, the **Online Supplementary Materials** and the **Online Supplementary Methods**.

#### Data availability

SRA accession numbers of sequencing data that has been deposited in NCBI, are listed per study in the **Online Supplementary Methods**.

#### Code availability

SMAP is available at https://gitlab.com/truttink/smap/ under the GNU Affero General Public License v3.0, and a detailed user manual is available at https://ngs-smap.readthedocs.io. Additional tools for downstream analysis of SMAP haplotype tables are available at https://gitlab.com/ybawin/smapapps and https://gitlab.com/ybawin/primer-design-gbs.

#### References (continued)

Baert J, *et al*. (2020). Breeding and genetics of two new amphiploid Festulolium synthetics with improved yield and digestibility. Biologia Plantarum 64:789-797 doi: 10.32615/bp.2020.138

Byrne S, *et al*. (2015). A synteny-based draft genome sequence of the forage grass *Lolium perenne*. The Plant Journal 84(4): 816-826. doi: 10.1111/tpj.13037

Chen HT, *et al*. (2020). Allele-aware chromosome-level genome assembly and efficient transgene-free genome editing for the autotetraploid cultivated alfalfa. Nature Communications 11(1):2494. doi: 10.1038/s41467-020-16338-x.

Danecek *et al*., (2011). The variant call format and VCFtools. Bioinformatics 27(15):2156-8. doi: 10.1093/bioinformatics/btr330.

De Bruyn C, *et al*. (2020). Establishment of CRISPR/Cas9 genome editing in Witloof (*Cichorium intybus* var. *foliosum*). Frontiers in genome editing: 24. doi: org./10.3389/fgeed.2020.604876.

Doyle JJ and Doyle JL. (1987). A rapid DNA isolation procedure for small quantities of fresh leaf tissue. Phytochemical Bulletin, 19(1), 11-15.

Elshire RJ, *et al*. (2011). A Robust, Simple Genotyping-by-Sequencing (GBS) Approach for High Diversity Species. PLoS ONE 6(5): e19379. doi: 10.1371/journal.pone.0019379.

Glaubitz JC, *et al*. (2014). TASSEL-GBS: A High Capacity Genotyping by Sequencing Analysis Pipeline. PLoS ONE 9(2): e90346. doi: 10.1371/journal.pone.0090346

Hapke A and Thiele D. (2016). GIbPSs: a toolkit for fast and accurate analyses of genotyping-by-sequencing data without a reference genome. Molecular Ecology Resources, 16(4): 979-990. doi: 10.1111/1755-0998.12510

Hardigan M, *et al*. (2016). Genome reduction uncovers a large dispensable genome and adaptive role for copy number variation in asexually propagated *Solanum tuberosum*. The Plant Cell 28(2): 388-405. doi: 10.1105/tpc.15.00538

Jiao WB and Schneeberger K. (2020). Chromosome-level assemblies of multiple Arabidopsis genomes reveal hotspots of rearrangements with altered evolutionary dynamics. Nature Communications 11(1): 989. doi: 10.1038/s41467-020-14779-y

Keep T, *et al*. (2020). High-Throughput Genome-Wide Genotyping To Optimize the Use of Natural Genetic Resources in the Grassland Species Perennial Ryegrass (*Lolium perenne* L.), G3 Genes|Genomes|Genetics 10(9): 3347–3364. doi: 10.1534/g3.120.401491

Kessner D, *et al*. (2013). Maximum likelihood estimation of frequencies of known haplotypes from pooled sequence data. Molecular Biology and Evolution 30(5): 1145–1158. doi: 10.1093/molbev/mst016

Li H. (2013). Aligning sequence reads, clone sequences and assembly contigs with BWA-MEM. arXiv preprint, 1303-3997.

Li H. (2018). Minimap2: pairwise alignment for nucleotide sequences. Bioinformatics 34(18): 3094-3100. doi: 10.1093/bioinformatics/bty191

Long Q, *et al*. (2011). PoolHap: Inferring haplotype frequencies from pooled samples by next generation sequencing. PloS ONE 6(1): e15292. doi: 10.1371/journal.pone.0015292

Lu F, *et al*. (2013). Switchgrass genomic diversity, ploidy, and evolution: novel insights from a network-based SNP discovery protocol. PLoS Genetics 9(1): e1003215. doi: 10.1371/journal.pgen.1003215

Manching H, *et al*. (2017). Phased Genotyping-by-Sequencing enhances analysis of genetic diversity and reveals divergent copy number variants in maize. G3 Genes|Genomes|Genetics, 7(7): 2161-2170. doi: 10.1534/g3.117.042036

Nagy I, *et al.* (2022). Chromosome-scale assembly and annotation of the perennial ryegrass genome. Submitted for publication.

O'Neil ST and Emrich SJ. (2012). Haplotype and minimum-chimerism consensus determination using short sequence data. BMC Genomics 13(Suppl 2): S4. doi: 10.1186/1471-2164-13-S2-S4

Page JT, *et al*. (2014). BamBam: genome sequence analysis tools for biologists. BMC Research Notes 7: 829. doi: 10.1186/1756-0500-7-829

Patterson M, *et al*. (2015). WhatsHap: Weighted Haplotype Assembly for Future-Generation Sequencing Reads. Journal of Computational Biology. 22(6): 498-509. doi: 10.1089/cmb.2014.0157

Scheet P and Stephens M. (2006). A fast and flexible statistical model for large-scale population genotype data: Applications to inferring missing genotypes and haplotypic phase. The American Journal of Human Genetics 78(4): 629-644. doi: 10.1086/502802

The Potato Genome Sequencing Consortium. (2011). Genome sequence and analysis of the tuber crop potato. Nature 475**,** 189–195. doi:10.1038/nature10158

Tinker NA, *et al*. (2016). Haplotag: Software for Haplotype-Based Genotyping-by-Sequencing Analysis. G3 Genes|Genomes|Genetics 6(4): 857-863. doi: 10.1534/g3.115.024596

Untergasser A., *et al*., (2012). Primer3—new capabilities and interfaces. Nucleic Acids Research 40(15), e115. doi: 10.1093/nar/gks596

Verwimp C, *et al.*, (2018). Temporal changes in genetic diversity and forage yield of perennial ryegrass in monoculture and in combination with red clover in swards. PLoS ONE 13(11): e0206571. doi: 10.1371/journal.pone.0206571

Wong T, *et al*. (2011). HaploJuice : accurate haplotype assembly from a pool of sequences with known relative concentrations. BMC Bioinformatics 19(1): 389. doi: 10.1186/s12859-018-2424-7

Zagordi O, *et al*. (2011). ShoRAH: estimating the genetic diversity of a mixed sample from next-generation sequencing data. BMC Bioinformatics 12(1): 119. doi: 10.1186/1471-2105-12-119

Zhang J, *et al*. (2014). PEAR: A fast and accurate Illumina Paired End reAd mergeR. Bioinformatics 30(5): 614–620. doi: 10.1093/bioinformatics/btt593
