## Supplementary material for "Stack Mapping Anchor Points (SMAP): a versatile suite of tools for read-backed haplotyping": Online_Supplemental_Methods

Methods and data analysis of demonstration case studies for:

**Analysis of PotatoMASH HiPlex data.** PotatoMASH (Potato Multi-Allele Scanning Haplotags) is a low cost, genome-scanning marker platform targeting 339 multi-allelic regions of the potato genome originally used to genotype 765 tetraploid potato lines comprising a training population derived from several crosses of multiple parents. Paired-end reads (unpublished results) were merged and mapped onto the *S. tuberosum* genome v4.04 (Hardigan *et al*., 2016) with BWA-MEM (Li, 2013). Out of 5,104 previously identified SNPs, 2,279 were selected based on minimum read depth 6 and minimum mapping quality 30 of polymorphisms across the potato lines. Using the primer binding site coordinates provided by the primer design software, we created a custom BED file with SMAPs defined as the first and last nucleotide of the amplicon, immediately internal to the primer pairs as the start and end points for haplotyping. *SMAP haplotype-sites* was run with parameters -mapping_orientation ignore -partial exclude --no_indels --discrete_calls dosage --frequency_interval_bounds 12.5 12.5 37.5 37.5 62.5 62.5 87.5 87.5 --dosage_filter 4 --min_read_count 20 --min_haplotype_frequency 5.

**Analysis of probe capture target enriched Shotgun sequencing of *Lolium perenne*.** Previously published probe capture target enriched Shotgun data (PE-91) of *L. perenne* diploid individuals (Veeckman *et al*., 2019) was re-analyzed with *SMAP haplotype-sites*. Out of 736 samples (BioProject PRJNA434356), 391 were selected based on library size (range 1.5-8 M reads per sample). Read data was mapped onto the *L. perenne* draft reference genome sequence (Byrne *et al*., 2015) with BWA-MEM (Li, 2013). Sliding frames were created for 253,125 previously identified bi-allelic SNPs (Veeckman *et al*., 2019) using the complementary python script provided as utility tools, with off-set 0, length range 10 to 80 bp in steps of 10 bp, frame distance 5. *SMAP haplotype-sites* was run with parameters -partial exclude -mapping_orientation ignore --q 30 --no_indels -c 10 -d 300 --min_distinct_haplotypes 2 --max_distinct_haplotypes 20 -f 5 --discrete_calls dosage --frequency_interval_bounds diploid --dosage_filter 2 --locus_correctness 80.

**Analysis of Structural Variants in Whole Genome Shotgun sequencing of *Oryza sativa*.** Previously published whole genome Shotgun data (WGS, PE-83/100/150) of *Oryza* diploid individuals and breakpoints of SVs (Kou *et al*., 2020) was re-analyzed with *SMAP haplotype-sites*. The VCF containing structural variant (SV) coordinates provided in the Supplementary Materials of Kou *et al*., (2020) was split into 72,930 DEL and 341,752 INV structural variants. Sliding frames were created using the complementary python script provided as *SMAP utility* tools, with off-set 1, length 3, frame distance 1. Out of 72,930 previously identified DELs, 71,973 non-overlapping 3bp sliding frames were selected for haplotyping. Out of 341,752 previously identified INVs, a random selection of 99,886 3bp non-overlapping sliding frames were selected for haplotyping. A saturation curve was created by computational subsampling (taking the top M reads of the fastq file at regular intervals, up to about 300M reads) of Nipponbare reads (SRA accession ERR2245546). In parallel, 269 unique samples (**Table S3**) were selected based on library size (range 5-300 M reads per sample), occasionally filling up gaps in the library size range coverage by computational subsampling of samples with large (over-saturated) library size. Read data was mapped onto the Nipponbare draft reference genome sequence (Kou *et al*., 2020) with BWA-MEM with default settings. *SMAP haplotype-sites* was run in parallel for the DEL and the INV SVs, with parameters: -mapping_orientation ignore -partial include --q 30 -d 300 --min_distinct_haplotypes 1 --max_distinct_haplotypes 20 -f 5 --discrete_calls dosage --frequency_interval_bounds 15 15 85 85 --dosage_filter 2, alternatively at read depth 15 (-c 15) or read depth 30 (-c 30).

| SRA | species | Bioproject | length | M reads |
|---|---|---|---|---|
| ERR619673 | Indica | PRJEB6180 | 166 | 5.6 |
| ERR619612 | Indica | PRJEB6180 | 166 | 5.6 |
| ERR625853 | Tropical japonica | PRJEB6180 | 166 | 5.6 |
| ERR620191 | Indica | PRJEB6180 | 166 | 5.6 |
| ERR625797 | Aus | PRJEB6180 | 166 | 5.6 |
| ERR625917 | Temperate japonica | PRJEB6180 | 166 | 5.7 |
| ERR624716 | Temperate japonica | PRJEB6180 | 166 | 5.7 |
| ERR623168 | Aus | PRJEB6180 | 166 | 5.9 |
| ERR625769 | Tropical japonica | PRJEB6180 | 166 | 5.9 |
| ERR606226 | Aus/boro | PRJEB6180 | 166 | 5.9 |
| ERR623201 | Tropical japonica | PRJEB6180 | 166 | 6.0 |
| ERR606338 | Tropical japonica | PRJEB6180 | 166 | 6.1 |
| ERR609009 | Temperate japonica | PRJEB6180 | 166 | 6.1 |
| ERR606506 | Aus/boro | PRJEB6180 | 166 | 6.1 |
| ERR621625 | Temperate japonica | PRJEB6180 | 166 | 6.2 |
| ERR619457 | Tropical japonica | PRJEB6180 | 166 | 6.2 |
| ERR624521 | Temperate japonica | PRJEB6180 | 166 | 6.5 |
| ERR622991 | Temperate japonica | PRJEB6180 | 166 | 6.6 |
| ERR623019 | Temperate japonica | PRJEB6180 | 166 | 6.6 |
| ERR623190 | Indica | PRJEB6180 | 166 | 6.7 |
| ERR611723 | Indica | PRJEB6180 | 166 | 6.7 |
| ERR606102 | Basmati/sadri | PRJEB6180 | 166 | 6.7 |
| ERR606436 | Aus/boro | PRJEB6180 | 166 | 6.8 |
| ERR606450 | Aus/boro | PRJEB6180 | 166 | 6.9 |
| ERR625102 | Indica | PRJEB6180 | 166 | 7.0 |
| ERR977887 | Indica | PRJEB6180 | 166 | 7.0 |
| ERR620519 | Temperate japonica | PRJEB6180 | 166 | 7.1 |
| ERR606520 | Aus/boro | PRJEB6180 | 166 | 7.1 |
| ERR622775 | Temperate japonica | PRJEB6180 | 166 | 7.1 |
| ERR630752 | Indica | PRJEB6180 | 166 | 7.4 |
| ERR978072 | Temperate japonica | PRJEB6180 | 166 | 7.4 |
| ERR607071 | Indica | PRJEB6180 | 166 | 7.4 |
| ERR622763 | Temperate japonica | PRJEB6180 | 166 | 7.4 |
| ERR624816 | Tropical japonica | PRJEB6180 | 166 | 7.5 |
| ERR636673 | Temperate japonica | PRJEB6180 | 166 | 7.5 |
| ERR631745 | Indica | PRJEB6180 | 166 | 7.5 |
| ERR611251 | Indica | PRJEB6180 | 166 | 7.6 |
| ERR623496 | Indica | PRJEB6180 | 166 | 7.7 |
| ERR622787 | Tropical japonica | PRJEB6180 | 166 | 7.9 |
| ERR636709 | Temperate japonica | PRJEB6180 | 166 | 8.0 |
| ERR630844 | Indica | PRJEB6180 | 166 | 8.1 |
| ERR611405 | Indica | PRJEB6180 | 166 | 8.1 |
| ERR618133 | Basmati/sadri | PRJEB6180 | 166 | 8.2 |
| ERR618339 | Aus/boro | PRJEB6180 | 166 | 8.2 |
| ERR611091 | Tropical japonica | PRJEB6180 | 166 | 8.3 |
| ERR624803 | Tropical japonica | PRJEB6180 | 166 | 8.4 |
| ERR623389 | Temperate japonica | PRJEB6180 | 166 | 8.4 |
| ERR622619 | Temperate japonica | PRJEB6180 | 166 | 8.4 |
| ERR607651 | Temperate japonica | PRJEB6180 | 166 | 8.5 |
| ERR630642 | Indica | PRJEB6180 | 166 | 8.5 |
| ERR608203 | Indica | PRJEB6180 | 166 | 8.5 |
| ERR611763 | Indica | PRJEB6180 | 166 | 8.5 |
| ERR618073 | Basmati/sadri | PRJEB6180 | 166 | 8.5 |
| ERR614201 | Basmati/sadri | PRJEB6180 | 166 | 8.6 |
| ERR630470 | Basmati/sadri | PRJEB6180 | 166 | 8.6 |
| ERR611485 | Indica | PRJEB6180 | 166 | 8.7 |
| ERR625050 | Temperate japonica | PRJEB6180 | 166 | 8.7 |
| ERR630714 | Temperate japonica | PRJEB6180 | 166 | 8.7 |
| ERR606560 | Basmati/sadri | PRJEB6180 | 166 | 8.7 |
| ERR631679 | Indica | PRJEB6180 | 166 | 8.8 |
| ERR631259 | Indica | PRJEB6180 | 166 | 8.8 |
| ERR636733 | Temperate japonica | PRJEB6180 | 166 | 8.9 |
| ERR630798 | Tropical japonica | PRJEB6180 | 166 | 8.9 |
| ERR608075 | Indica | PRJEB6180 | 166 | 9.1 |
| ERR611385 | Indica | PRJEB6180 | 166 | 9.1 |
| ERR636979 | Temperate japonica | PRJEB6180 | 166 | 9.1 |
| ERR608257 | Indica | PRJEB6180 | 166 | 9.3 |
| ERR606214 | Aus/boro | PRJEB6180 | 166 | 9.3 |

| SRA | species | Bioproject | length | M reads |
|---|---|---|---|---|
| ERR637009 | Temperate japonica | PRJEB6180 | 166 | 9.6 |
| ERR611637 | Temperate japonica | PRJEB6180 | 166 | 9.6 |
| ERR607817 | Indica | PRJEB6180 | 166 | 9.7 |
| ERR631725 | Indica | PRJEB6180 | 166 | 9.8 |
| ERR618717 | Tropical japonica | PRJEB6180 | 166 | 9.9 |
| ERR631613 | Indica | PRJEB6180 | 166 | 9.9 |
| ERR618585 | Tropical japonica | PRJEB6180 | 166 | 10.1 |
| ERR631593 | Indica | PRJEB6180 | 166 | 10.1 |
| ERR605976 | Tropical japonica | PRJEB6180 | 166 | 10.1 |
| ERR617903 | Tropical japonica | PRJEB6180 | 166 | 10.2 |
| ERR607899 | Tropical japonica | PRJEB6180 | 166 | 10.2 |
| ERR624985 | Temperate japonica | PRJEB6180 | 166 | 10.2 |
| ERR630464 | Indica | PRJEB6180 | 166 | 10.2 |
| ERR608185 | Tropical japonica | PRJEB6180 | 166 | 10.3 |
| ERR605635 | Tropical japonica | PRJEB6180 | 166 | 10.3 |
| ERR615780 | Indica | PRJEB6180 | 166 | 10.3 |
| ERR630694 | Temperate japonica | PRJEB6180 | 166 | 10.3 |
| ERR621937 | Temperate japonica | PRJEB6180 | 166 | 10.3 |
| ERR608907 | Temperate japonica | PRJEB6180 | 166 | 10.6 |
| ERR630476 | Basmati/sadri | PRJEB6180 | 166 | 10.7 |
| ERR637051 | Temperate japonica | PRJEB6180 | 166 | 10.7 |
| ERR605509 | Aus/boro | PRJEB6180 | 166 | 10.8 |
| ERR618163 | Tropical japonica | PRJEB6180 | 166 | 10.8 |
| ERR612969 | Aus/boro | PRJEB6180 | 166 | 10.9 |
| ERR631080 | Indica | PRJEB6180 | 166 | 10.9 |
| ERR630500 | Basmati/sadri | PRJEB6180 | 166 | 10.9 |
| ERR608683 | Temperate japonica | PRJEB6180 | 166 | 11.0 |
| ERR607615 | Tropical japonica | PRJEB6180 | 166 | 11.0 |
| ERR616652 | Basmati/sadri | PRJEB6180 | 166 | 11.1 |
| ERR605485 | Tropical japonica | PRJEB6180 | 166 | 11.1 |
| ERR622583 | Temperate japonica | PRJEB6180 | 166 | 11.1 |
| ERR611975 | Indica | PRJEB6180 | 166 | 11.2 |
| ERR616060 | Indica | PRJEB6180 | 166 | 11.2 |
| ERR605701 | Tropical japonica | PRJEB6180 | 166 | 11.2 |
| ERR618181 | Indica | PRJEB6180 | 166 | 11.4 |
| ERR607995 | Tropical japonica | PRJEB6180 | 166 | 11.6 |
| ERR618321 | Tropical japonica | PRJEB6180 | 166 | 11.6 |
| ERR612535 | Indica | PRJEB6180 | 166 | 11.7 |
| ERR612119 | Tropical japonica | PRJEB6180 | 166 | 11.8 |
| ERR618187 | Indica | PRJEB6180 | 166 | 11.8 |
| ERR635284 | Basmati/sadri | PRJEB6180 | 166 | 11.8 |
| ERR636955 | Temperate japonica | PRJEB6180 | 166 | 11.8 |
| ERR626819 | Tropical japonica | PRJEB6180 | 166 | 11.9 |
| ERR608287 | Temperate japonica | PRJEB6180 | 166 | 11.9 |
| ERR626441 | Indica | PRJEB6180 | 166 | 12.1 |
| ERR605587 | Indica | PRJEB6180 | 166 | 12.2 |
| ERR613890 | Tropical japonica | PRJEB6180 | 166 | 12.3 |
| ERR630864 | Indica | PRJEB6180 | 166 | 12.3 |
| ERR613683 | Temperate japonica | PRJEB6180 | 166 | 12.5 |
| ERR608111 | Indica | PRJEB6180 | 166 | 12.5 |
| ERR605429 | Temperate japonica | PRJEB6180 | 166 | 12.6 |
| ERR617649 | Tropical japonica | PRJEB6180 | 166 | 12.6 |
| ERR613990 | Tropical japonica | PRJEB6180 | 166 | 12.7 |
| ERR617277 | Tropical japonica | PRJEB6180 | 166 | 12.9 |
| ERR616967 | Indica | PRJEB6180 | 166 | 13.3 |
| ERR614088 | Indica | PRJEB6180 | 166 | 13.4 |
| ERR615033 | Indica | PRJEB6180 | 166 | 13.6 |
| ERR608775 | Tropical japonica | PRJEB6180 | 166 | 13.7 |
| ERR613437 | Tropical japonica | PRJEB6180 | 166 | 13.8 |
| ERR629958 | Indica | PRJEB6180 | 166 | 14.0 |
| ERR613866 | Tropical japonica | PRJEB6180 | 166 | 14.3 |
| ERR626734 | Indica | PRJEB6180 | 166 | 14.4 |
| ERR613533 | Indica | PRJEB6180 | 166 | 14.7 |
| ERR613878 | Tropical japonica | PRJEB6180 | 166 | 14.8 |
| ERR612915 | Indica | PRJEB6180 | 166 | 15.0 |
| ERR613017 | Indica | PRJEB6180 | 166 | 15.6 |
| ERR613407 | Tropical japonica | PRJEB6180 | 166 | 15.6 |
| ERR613011 | Indica | PRJEB6180 | 166 | 16.5 |

| SRA | species | Bioproject | length | M reads |
|---|---|---|---|---|
| SRR3180882 | Oryza rufipogon | PRJNA312733 | 180 | 209.7 |
| DRR001186 | Wild rice O. rufipogo | PRJDB2009 | 202 | 95.2 |
| DRR001188 | Wild rice O. rufipogo | PRJDB2009 | 202 | 84.8 |
| DRR001183 | Wild rice O. rufipogo | PRJDB2009 | 202 | 100.4 |
| DRR001185 | Wild rice O. rufipogo | PRJDB2009 | 202 | 112.5 |
| DRR001189 | Wild rice O. rufipogo | PRJDB2009 | 202 | 93.8 |
| DRR001184 | Wild rice O. rufipogo | PRJDB2009 | 202 | 96.9 |
| DRR001187 | Wild rice O. rufipogo | PRJDB2009 | 202 | 4.9 |
| DRR001190 | Wild rice O. rufipogo | PRJDB2009 | 202 | 92.4 |
| SRR5536054 | Oryza rufipogon | PRJNA382258 | 202 | 114.9 |
| SRR5536055 | Oryza rufipogon | PRJNA382258 | 202 | 128.8 |
| SRR1450140 | Oryza sativa f. sponta | PRJNA48107 | 200 | 47.6 |
| SRR1450141 | Oryza sativa f. sponta | PRJNA48107 | 200 | 38.2 |
| SRR1707282 | Oryza sativa f. sponta | PRJNA264484 | 181 | 31.3 |
| SRR1743122 | Oryza sativa f. sponta | PRJNA271253 | 200 | 26.2 |
| SRR1743094 | Oryza sativa Indica Gr | PRJNA271253 | 200 | 66.4 |
| ERR2240128 | Oryza sativa | PRJEB19404 | 200 | 261.0 |
| ERR2245546 | Oryza sativa Japonica | PRJEB19404 | 200 | 301.7 |
| SRR1450058* | Oryza meridionalis | PRJNA48433 | 200 | 57.9 |
| ERR2241057* | Oryza sativa | PRJEB19404 | 200 | 69.8 |
| ERR2240124* | Oryza sativa | PRJEB19404 | 200 | 77.3 |
| ERR2242621* | Oryza sativa | PRJEB19404 | 200 | 79.8 |
| SRR3234372 | Oryza sativa indica | PRJNA276972 | 202 | 255.2 |
| SRR3234369 | Oryza sativa Indica Gr | PRJNA276972 | 202 | 284.9 |
| DRR088694 | Oryza rufipogon | PRJDB5512 | 300 | 23.8 |
| DRR088687 | Oryza rufipogon | PRJDB5512 | 300 | 24.0 |
| DRR088689 | Oryza rufipogon | PRJDB5512 | 300 | 24.5 |
| DRR088688 | Oryza rufipogon | PRJDB5512 | 300 | 25.0 |
| SRR8324888 | Oryza rufipogon | PRJNA260762 | 300 | 25.6 |
| SRR8324908 | Oryza rufipogon | PRJNA260762 | 300 | 26.2 |
| DRR088677 | Oryza rufipogon | PRJDB5512 | 300 | 26.6 |
| DRR088676 | Oryza rufipogon | PRJDB5512 | 300 | 27.3 |
| DRR088693 | Oryza rufipogon | PRJDB5512 | 300 | 27.9 |
| SRR8324913 | Oryza rufipogon | PRJNA260762 | 300 | 27.9 |
| SRR8324877 | Oryza rufipogon | PRJNA260762 | 300 | 28.4 |
| SRR8324907 | Oryza rufipogon | PRJNA260762 | 300 | 28.6 |
| SRR8324916 | Oryza rufipogon | PRJNA260762 | 300 | 28.7 |
| SRR8324919 | Oryza rufipogon | PRJNA260762 | 300 | 29.2 |
| DRR088690 | Oryza rufipogon | PRJDB5512 | 300 | 29.2 |
| DRR058041 | Oryza longistaminata | PRJDB4705 | 300 | 29.3 |
| DRR088686 | Oryza rufipogon | PRJDB5512 | 300 | 29.8 |
| DRR088683 | Oryza rufipogon | PRJDB5512 | 300 | 30.2 |
| SRR8324860 | Oryza rufipogon | PRJNA260762 | 300 | 30.4 |
| DRR088680 | Oryza rufipogon | PRJDB5512 | 300 | 30.5 |
| SRR8324880 | Oryza rufipogon | PRJNA260762 | 300 | 30.5 |
| SRR8324882 | Oryza rufipogon | PRJNA260762 | 300 | 31.0 |
| SRR8324885 | Oryza rufipogon | PRJNA260762 | 300 | 31.2 |
| SRR8324857 | Oryza rufipogon | PRJNA260762 | 300 | 31.2 |
| SRR8324911 | Oryza rufipogon | PRJNA260762 | 300 | 31.3 |
| DRR088692 | Oryza rufipogon | PRJDB5512 | 300 | 31.4 |
| SRR8324915 | Oryza rufipogon | PRJNA260762 | 300 | 31.4 |
| DRR088691 | Oryza rufipogon | PRJDB5512 | 300 | 32.0 |
| SRR8324887 | Oryza rufipogon | PRJNA260762 | 300 | 32.0 |
| SRR8324891 | Oryza rufipogon | PRJNA260762 | 300 | 32.1 |
| SRR8324909 | Oryza rufipogon | PRJNA260762 | 300 | 32.2 |
| DRR088684 | Oryza rufipogon | PRJDB5512 | 300 | 32.4 |
| SRR8324886 | Oryza rufipogon | PRJNA260762 | 300 | 32.4 |
| SRR8324910 | Oryza rufipogon | PRJNA260762 | 300 | 32.6 |
| SRR8324918 | Oryza rufipogon | PRJNA260762 | 300 | 32.6 |
| SRR8324892 | Oryza rufipogon | PRJNA260762 | 300 | 33.1 |
| DRR088682 | Oryza rufipogon | PRJDB5512 | 300 | 33.1 |
| DRR088675 | Oryza rufipogon | PRJDB5512 | 300 | 33.2 |
| SRR8324884 | Oryza rufipogon | PRJNA260762 | 300 | 33.5 |
| SRR8324878 | Oryza rufipogon | PRJNA260762 | 300 | 33.8 |
| SRR8324861 | Oryza rufipogon | PRJNA260762 | 300 | 34.0 |
| SRR8324879 | Oryza rufipogon | PRJNA260762 | 300 | 34.2 |
| SRR8324895 | Oryza rufipogon | PRJNA260762 | 300 | 34.6 |

| SRA | species | Bioproject | length | M reads |
|---|---|---|---|---|
| SRR8324859 | Oryza rufipogon | PRJNA260762 | 300 | 34.9 |
| DRR088685 | Oryza rufipogon | PRJDB5512 | 300 | 35.0 |
| SRR8324881 | Oryza rufipogon | PRJNA260762 | 300 | 35.0 |
| SRR8324883 | Oryza rufipogon | PRJNA260762 | 300 | 35.4 |
| SRR8324912 | Oryza rufipogon | PRJNA260762 | 300 | 35.5 |
| SRR8324862 | Oryza rufipogon | PRJNA260762 | 300 | 36.3 |
| SRR8324890 | Oryza rufipogon | PRJNA260762 | 300 | 36.4 |
| SRR8324864 | Oryza rufipogon | PRJNA260762 | 300 | 36.4 |
| SRR8324889 | Oryza rufipogon | PRJNA260762 | 300 | 36.4 |
| SRR8324896 | Oryza rufipogon | PRJNA260762 | 300 | 36.5 |
| SRR8324920 | Oryza rufipogon | PRJNA260762 | 300 | 37.2 |
| SRR8324914 | Oryza rufipogon | PRJNA260762 | 300 | 37.2 |
| DRR088681 | Oryza rufipogon | PRJDB5512 | 300 | 38.4 |
| SRR8324863 | Oryza rufipogon | PRJNA260762 | 300 | 38.5 |
| SRR8324894 | Oryza rufipogon | PRJNA260762 | 300 | 39.4 |
| ERR2245531 | Oryza sativa | PRJEB19404 | 300 | 48.0 |
| ERR2241059 | Oryza sativa | PRJEB19404 | 300 | 50.5 |
| ERR2245530 | Oryza sativa | PRJEB19404 | 300 | 59.9 |
| ERR2245541 | Oryza sativa | PRJEB19404 | 300 | 62.7 |
| ERR2245545 | Oryza sativa | PRJEB19404 | 300 | 63.3 |
| ERR2245532 | Oryza sativa | PRJEB19404 | 300 | 67.8 |
| ERR2245515 | Oryza sativa | PRJEB19404 | 300 | 106.8 |
| ERR2245544 | Oryza sativa | PRJEB19404 | 300 | 113.5 |
| ERR2245536 | Oryza sativa | PRJEB19404 | 300 | 118.6 |
| ERR2245519 | Oryza sativa | PRJEB19404 | 300 | 120.6 |
| ERR2245527 | Oryza sativa | PRJEB19404 | 300 | 126.7 |
| ERR2245556 | Oryza rufipogon | PRJEB19404 | 300 | 130.8 |
| ERR2245524 | Oryza sativa | PRJEB19404 | 300 | 145.1 |
| ERR2245533 | Oryza sativa | PRJEB19404 | 300 | 146.4 |
| ERR2245534 | Oryza sativa | PRJEB19404 | 300 | 148.3 |
| ERR2245523 | Oryza sativa | PRJEB19404 | 300 | 149.3 |
| ERR2245517 | Oryza sativa | PRJEB19404 | 300 | 152.5 |
| ERR2245550 | Oryza rufipogon | PRJEB19404 | 300 | 162.0 |
| ERR2245513 | Oryza sativa | PRJEB19404 | 300 | 172.1 |
| ERR2245551 | Oryza rufipogon | PRJEB19404 | 300 | 172.7 |
| ERR2242625 | Oryza sativa | PRJEB19404 | 300 | 179.1 |
| ERR2245516 | Oryza sativa | PRJEB19404 | 300 | 185.9 |
| ERR2242624 | Oryza sativa | PRJEB19404 | 300 | 186.6 |
| ERR2245539 | Oryza sativa | PRJEB19404 | 300 | 199.0 |
| ERR2245549 | Oryza rufipogon | PRJEB19404 | 300 | 204.3 |
| ERR2245520 | Oryza sativa | PRJEB19404 | 300 | 205.4 |
| ERR2245548 | Oryza rufipogon | PRJEB19404 | 300 | 206.4 |
| ERR2245525 | Oryza sativa | PRJEB19404 | 300 | 207.3 |
| ERR2245512 | Oryza sativa | PRJEB19404 | 300 | 209.2 |
| ERR2240125 | Oryza rufipogon | PRJEB19404 | 300 | 211.1 |
| ERR2240126 | Oryza rufipogon | PRJEB19404 | 300 | 214.9 |
| ERR2241058 | Oryza sativa | PRJEB19404 | 300 | 224.2 |
| ERR2245552 | Oryza rufipogon | PRJEB19404 | 300 | 226.8 |
| ERR2245529 | Oryza sativa | PRJEB19404 | 300 | 238.9 |
| ERR2245554 | Oryza rufipogon | PRJEB19404 | 300 | 248.6 |
| ERR2241055 | Oryza sativa | PRJEB19404 | 300 | 250.7 |
| ERR2245557 | Oryza rufipogon | PRJEB19404 | 300 | 256.0 |
| ERR2245528 | Oryza sativa | PRJEB19404 | 300 | 273.0 |
| ERR2240127 | Oryza sativa | PRJEB19404 | 300 | 286.7 |
| ERR2241056 | Oryza sativa | PRJEB19404 | 300 | 308.1 |
| ERR2242626* | Oryza sativa | PRJEB19404 | 300 | 20.7 |
| ERR2245535* | Oryza sativa | PRJEB19404 | 300 | 30.9 |
| ERR2245537* | Oryza sativa | PRJEB19404 | 300 | 34.6 |
| ERR2245538* | Oryza sativa | PRJEB19404 | 300 | 93.1 |
| ERR2245542* | Oryza sativa | PRJEB19404 | 300 | 31.1 |
| ERR2241060* | Oryza sativa | PRJEB19404 | 300 | 39.5 |
| ERR2242623* | Oryza sativa | PRJEB19404 | 300 | 56.7 |
| SRR8241155 | Oryza sativa L. ssp. indica | PRJNA482013 | 300 | 124.9 |
| SRR5011847 | Oryza sativa Indica Group | PRJNA318714 | 300 | 177.0 |
| SRR3422892 | Oryza sativa Indica Group | PRJNA318714 | 300 | 187.0 |
| SRR5880534 | early-matured japonica (Geng) | PRJNA396422 | 300 | 125.2 |

**Table S3 | Selection of 269 WGS datasets of *Oryza sativa* for haplotype calling of structural variants.** Table lists SRA accession number, species, Bioproject number, total read length (*e.g.* PE-150 = 300 bp), and number of reads mapped (in M). Samples with over-saturated library size in SRA that were computational subsampled to fill gaps in the library size range coverage are indicated by an *.

**Analysis of PacBio long range whole genome resequencing data of *Arabidopsis thaliana*.** Previously published PacBio data (Jiao and Schneeberger, 2020), of seven *Arabidopsis* diploid ecotypes (An-1, C24, Cvi, Eri, Kyo, Ler, Sha) was re-analyzed with *SMAP haplotype-sites*. Read data was downloaded from SRA (Bioproject PRJEB31147) and mapped onto the Tair10 *Arabidopsis* reference genome sequence (www.arabidopsis.org/) with Minimap2 (Li, 2018). SNPs were retrieved from www.1001genomes.org/accessions.html, and 1,223,090 bi-allelic SNPs that were polymorphic in six out of seven ecotypes (An-1, Cvi, Eri, Kyo, Ler, Sha; no SNP data was available for C24) were selected. Sliding frames were created using the complementary python script provided as *SMAP utility* tools on GitLab, with off-set 0, length range 250 bp to 3000 bp in steps of 250 bp, frame distance 0. *SMAP haplotype-sites* was run with parameters -mapping_orientation ignore -partial exclude --q 30 --no_indels -c 10 -d 300 --min_distinct_haplotypes 2 --max_distinct_haplotypes 20 -f 5 --discrete_calls dosage --frequency_interval_bounds 30 30 70 70 --dosage_filter 2 --locus_correctness 100.

**GBS: Analysis of SNPs affecting restriction sites in *L. perenne*.** We analyzed the sequence context of SNPs identified in a broad collection of *L. perenne* genotypes to quantify the expected frequency with which restriction sites in the reference genome sequence are affected by genome sequence diversity within the genepool – leading to missing data during genotype calling across sets of individuals or pools. Targeted resequencing with a probe capture based method (hence independent of the distribution of restriction sites in each sample) in 736 genotypes across 2.3 Mb of genome sequence yielded 253,125 bi-allelic SNPs (BioProject PRJNA434356 and Supplementary materials of Veeckman *et al*., 2019). The reference genome sequence context per SNP was extracted to interrogate if the switch from the reference nucleotide to the alternative nucleotide leads to the loss or gain of a restriction site. In addition, if the loss of a restriction site simultaneously leads to the creation of a novel restriction site, this leads to a small shift in the restriction site position, thus creating an additional SMAP (**Fig. 3f**). SNPs were classified based on their relative abundance (expressed in minor allele frequency (MAF) in the collection of 736 genotypes. We found that a substantial number of restriction sites in the *L. perenne* reference genome sequence (Byrne *et al*., 2015) are affected upon substitution of the reference nucleotide to the alternative nucleotide (**Supplementary Table S2**). For instance, 129 (14.5%) of the 1218 *Pst*I restriction sites (contained in the 2.3 Mb resequenced genome fraction) are lost, and 177 (10.6%) restriction sites are gained due to moderate frequency SNPs (MAF>5%), while another 103 (8.5%) and 95 (7.8%) restriction sites are lost and gained, respectively, due to low frequency SNPs (MAF 1%-5%).These numbers are consistent across different enzymes, and as expected, more GBS loci are affected when performing GBS with restriction enzymes with shorter target recognition sequence (*e.g.* four-cutters). Taken together, these analyses show that in heterogeneous populations of heterozygous individuals in outcrossing species, up to one-quarter or one-third of the GBS stack positions may be affected by the loss or gain of restriction sites, albeit by relatively low MAF SNPs. These cause the presence/absence of loci per sample and explain the relatively high fraction of individual-private and accession-private loci, as identified by *SMAP delineate* (**Fig. 3d,e**).

**GBS: *in silico* prediction of loci on the reference sequence versus *SMAP delineate*.** To further illustrate the importance of this effect on real data, we mapped *Pst*I-GBS data of 48 *L. perenne* individuals (BioProject PRJNA812749) onto the reference genome (Byrne *et al*., 2015) with BWA-MEM, and determined the GBS StackCluster coordinates with *SMAP delineate*. Next, we compared those GBS loci with predicted GBS locus positions in the reference genome based on an *in-silico* digest and computational fragment size selection (50-500 bp). The comparison showed that about one-third of the loci covered by mapped reads did not overlap with predicted GBS loci in the reference genome, and, *vice versa*, up to one-third of the predicted GBS loci was not covered by reads in any of the 48 individuals. From the loci that are found outside the *in silico* predicted mapping sites, many loci typically only contain read data from individual genotypes (individual-private loci, **Fig. 3d**), showing that polymorphisms in restriction sites create additional and unique stacks in individual genotypes. This analysis revealed that an *in silico* digest of the reference genome in a highly polymorphic sample set, is a poor predictor of the GBS stack locations, causing loss of data for algorithms that rely on a predefined BED file delineating GBS loci (*e.g.* haplotyping by LocHap-GBS; Manching *et al*., 2017). Furthermore, quantification of the fraction of StackClusters that are affected by insertions, deletions, soft-clipping and hard-clipping (**Fig. 3g**), further shows that stacks frequently do not align entirely to the expected regions (i.e. directly flanking the restriction sites) within a given locus.

**GBS: optimization of restriction enzymes for *M. sativa*.** A pilot study to optimize the choice of restriction enzymes in *M. sativa* was previously published (Julier *et al*., 2021), and read data were re-analyzed to Illustrate the use of *SMAP delineate*. Reads were mapped onto the reference genome (Chen *et al*., 2020) with BWA-MEM with default parameters. GBS loci were identified with *SMAP delineate* with parameters -mapping_orientation ignore -p 34 --min_cluster_length 40 --max_cluster_length 300 --min_stack_depth 10 --max_stack_depth 3000 --min_cluster_depth 30 --max_cluster_depth 3000 --max_stack_number 20 --min_stack_depth_fraction 5 --max_smap_number 20, --completeness 1 --plot all.

**GBS: Analysis of Pool-Seq data in natural populations of *L. perenne*.** Previously published Pool-GBS data (SE-86; BioProject PRJNA445949) of 552 *L. perenne* genebank accessions (Blanco-Pastor *et al*., 2019, Keep *et al*., 2020) was re-analyzed with *SMAP delineate*. Read data was mapped onto the *L. perenne* draft reference genome sequence (Byrne *et al*., 2015) with BWA-MEM. GBS loci were identified with *SMAP delineate* with parameters: -mapping_orientation stranded -p 24 --name lolium_GLS_552 -f 50 -g 120 --min_stack_depth 2 --max_stack_depth 1500 --min_cluster_depth 30 --max_cluster_depth 3000 --max_stack_number 20 --min_stack_depth_fraction 5 --max_smap_number 10. *SMAP haplotype-sites* was run with parameters: -mapping_orientation stranded -partial include --q 30 --no_indels -c 30 -d 300 --min_distinct_haplotypes 2 --max_distinct_haplotypes 20 -f 5 -m 1.

**GBS/HiPlex: Iterative cycles of SMAP for identification of interspecific hybrids in the *Festulolium* complex.** *Pst*I-GBS data (PE-150; BioProject PRJNA812869) of 11 *L. perenne,* 12 *L. multiflorum* and 14 *F. pratensis* parental lines were generated following the protocol of Verwimp *et al*. (2018). Read data were prepared for read mapping by *GBprocesS* and mapped onto the *L. perenne* reference genome sequence (Nagy *et al*., submitted for publication, https://ryegrassgenome.ghpc.au.dk/) with BWA-MEM. GBS loci were identified with *SMAP delineate* with parameters: -mapping_orientation ignore -p 8 --plot all --plot_type png --name 2n_ind_GBS-PE -f 90 -g 300 --min_stack_depth 2 --max_stack_depth 500 --min_cluster_depth 10 --max_cluster_depth 1500 --max_stack_number 2 --min_stack_depth_fraction 10 --completeness 80 --max_smap_number 2. SNP calling and filtering was performed with VCFtools (Danecek *et al*., 2011) and GATK and only polymorphic bi-allelic SNPs were used to run *SMAP haplotype-sites* with parameters: -mapping_orientation ignore -partial include --no_indels --min_read_count 10 --min_distinct_haplotypes 2 -f 5 -p 16 --discrete_calls dosage -i 10 10 90 90 -z 2. The *SMAP utility* tool *SMAPapp-Matrix.py* (https://gitlab.com/ybawin/smapapps) was used to create a matrix showing the number of loci with only unique haplotypes in each pair of parental lines with parameters: python3 SMAPapp-Matrix.py --table haplotypes_c10_f5_m0_discrete_calls_filtered.tsv -lic Unique --partial False --proportion_informative_loci False --print_locus_information All -rc 0.10 --plot_format png --annotate_matrix_plots. HiPlex primer design within GBS loci with the highest locus information content was performed with Primer3 (Untergasser *et al*., 2012) implemented in the *Primer_design.py* script (https://gitlab.com/ybawin/primer-design-gbs) with parameters --vcf snps_GBS_polymorph_biallelic.vcf --reference Lolium_2.6.1.fa --regions regions_best_locus_information_content.bed --maximum_amplicon_size 120. The discriminatory power of the designed primer set was checked *in silico* by running *SMAP haplotype-sites* on the selected HiPlex loci in the GBS data with parameters: SMAP_haplotype_sites_primer_set_272.bed snps_GBS_polymorph_biallelic.vcf -mapping_orientation ignore -partial include --no_indels --min_read_count 10 --min_distinct_haplotypes 2 -f 5 -p 16 --plot_type png --discrete_calls dosage -i 10 10 90 90 -z 2, followed by running *SMAPapp-Matrix.py* again with the same parameters as above but with the haplotypes table based on the selected HiPlex primer loci. Iterative cycles of *Primer_design.py*, *SMAP haplotype-sites* and *SMAPapp-Matrix.py* were run to *in silico* design a primer set with increasingly higher discriminatory power and finally yielded a set of 272 loci that could discriminate all parental lines. Interspecific *Festuca-Lolium* hybrids were made according to Baert *et al*., (2020). HiPlex data (PE-150, BioProject PRJNA812869) of 272 amplicons in 11 *L. perenne,* 12 *L. multiflorum* and 14 *F. pratensis* parental lines and 85 progeny plants were used to identify interspecific hybrids in the *Festulolium* complex by mapping the reads onto the *L. perenne* reference genome sequence with BWA-MEM. SNP calling and filtering was performed with VCFtools (Danecek *et al*., 2011) and *GATK* and only polymorphic bi-allelic SNPs were used to run *SMAP haplotype-sites* with parameters: SMAP_haplotype_sites_primer_set_272.bed snps_HiPlex_polymorph_biallelic.vcf -mapping_orientation ignore -partial exclude --no_indels --min_read_count 10 -f 5 -p 16 --plot all --discrete_calls dominant -i 10. The *SMAP utility* tool *SMAPapp-Matrix.py* was used to compare potential hybrids with their respective parents with parameters: -i input_directory -o output_directory -n sample_names_parents-hybrids.txt --print_sample_information All --print_locus_information All -p 24 --distance --distance_method Inversed -lic Shared --partial --annotate_matrix_plots --title_fontsize 22 --label_fontsize 10 --tick_fontsize 6 -sc 10 -lc 0.1 --plot_format pdf --legend_fontsize 10 --proportion_informative_loci True -u 'parents-hybrids_PartiallyShared'. The partial similarity score is the proportion of loci with partially shared haplotypes, the highest proportion corresponds to the respective parents.

**Multiplex CRISPR/Cas genome editing in potato.** gRNAs (Geneious Prime 2.0) and HiPlex amplicon primers (Primer3) were designed for 88 candidate genes using the annotated reference genome of *Solanum tuberosum* (The Potato Genome Sequencing Consortium, 2011). A HiPlex assay with 222 amplicons was tested on wild type genomic DNA of Bintje and Spunta, and loci for genome editing/targeted resequencing were selected based on good amplification per locus, unambiguous detection of four alleles in tetraploid wild type background per cultivar, and possibility to design gRNAs targeting both Bintje and Spunta alleles. In this study, 44 gRNAs targeting 33 genes were selected for further analysis, cloned in gRNA expression vectors (one gRNA per vector), and divided into four pools with 11 gRNA vectors for co-transfection. Protoplast isolation, cloning of Cas9 and gRNA expression vectors, PEG co-transfection, and plant regeneration was performed according to the CRISPR/Cas genome editing protocol previously described by De Bruyn *et al*., (2020). DNA was isolated from regenerated calli (either pools of five calli or single calli) using a modified CTAB protocol according to Doyle and Doyle (1987). HiPlex reads were mapped to the candidate gene sequences extracted from the *S. tuberosum* reference genome, and borders (10 bp) delineating windows were custom defined based on HiPlex primer binding positions. *SMAP haplotype-window* was run with parameters: -p 24 -min_distinct_haplotypes 0 -f 5 -c 30 -m 2, without discrete genotype call options activated because the sample set combined pool-Seq HiPlex data and tetraploid individual calli.

**Affiliations**

[1]Flanders Research Institute for Agriculture, Fisheries and Food, Plant Sciences Unit, Caritasstraat 39, B-9090, Melle, Belgium, [2]Ghent University, Department of Applied Mathematics, Computer Science and Statistics, Krijgslaan 281 S9, B-9000, Ghent, Belgium, [3]INRAE, URP3F, F-86600 Lusignan, France, [4]Teagasc, Crop Science Department, Oak Park, Carlow R93 XE12, Ireland. Correspondence should be addressed to T.R..

**Methods**

Methods, including a detailed user manual explaining the features and optional parameters of the code, statements of data availability and any associated accession codes and references, are available in the **Online manual**, the **Online Supplementary Materials** and the **Online Supplementary Methods**.

**Data availability**

SRA accession numbers of sequencing data that has been deposited in NCBI, are listed per study in the **Online Supplementary Methods**.

**Code availability**

SMAP is available at https://gitlab.com/truttink/smap/ under the GNU Affero General Public License v3.0, and a detailed user manual is available at https://ngs-smap.readthedocs.io. Additional tools for downstream analysis of SMAP haplotype tables are available at https://gitlab.com/ybawin/smapapps and https://gitlab.com/ybawin/primer-design-gbs.

**References (continued)**

Baert J, *et al*. (2020). Breeding and genetics of two new amphiploid Festulolium synthetics with improved yield and digestibility. Biologia Plantarum 64:789-797 doi: 10.32615/bp.2020.138

Byrne S, *et al*. (2015). A synteny-based draft genome sequence of the forage grass *Lolium perenne*. The Plant Journal 84(4): 816-826. doi: 10.1111/tpj.13037

Chen HT, *et al*. (2020). Allele-aware chromosome-level genome assembly and efficient transgene-free genome editing for the autotetraploid cultivated alfalfa. Nature Communications 11(1):2494. doi: 10.1038/s41467-020-16338-x.

Danecek *et al*., (2011). The variant call format and VCFtools. Bioinformatics 27(15):2156-8. doi: 10.1093/bioinformatics/btr330.

De Bruyn C, *et al*. (2020). Establishment of CRISPR/Cas9 genome editing in Witloof (*Cichorium intybus* var. *foliosum*). Frontiers in genome editing: 24. doi: org./10.3389/fgeed.2020.604876.

Doyle JJ and Doyle JL. (1987). A rapid DNA isolation procedure for small quantities of fresh leaf tissue. Phytochemical Bulletin, 19(1), 11-15.

Elshire RJ, *et al*. (2011). A Robust, Simple Genotyping-by-Sequencing (GBS) Approach for High Diversity Species. PLoS ONE 6(5): e19379. doi: 10.1371/journal.pone.0019379.

Glaubitz JC, *et al*. (2014). TASSEL-GBS: A High Capacity Genotyping by Sequencing Analysis Pipeline. PLoS ONE 9(2): e90346. doi: 10.1371/journal.pone.0090346

Hapke A and Thiele D. (2016). GIbPSs: a toolkit for fast and accurate analyses of genotyping-by-sequencing data without a reference genome. Molecular Ecology Resources, 16(4): 979-990. doi: 10.1111/1755-0998.12510

Hardigan M, *et al*. (2016). Genome reduction uncovers a large dispensable genome and adaptive role for copy number variation in asexually propagated *Solanum tuberosum*. The Plant Cell 28(2): 388-405. doi: 10.1105/tpc.15.00538

Jiao WB and Schneeberger K. (2020). Chromosome-level assemblies of multiple Arabidopsis genomes reveal hotspots of rearrangements with altered evolutionary dynamics. Nature Communications 11(1): 989. doi: 10.1038/s41467-020-14779-y

Keep T, *et al*. (2020). High-Throughput Genome-Wide Genotyping To Optimize the Use of Natural Genetic Resources in the Grassland Species Perennial Ryegrass (*Lolium perenne* L.), G3 Genes|Genomes|Genetics 10(9): 3347–3364. doi: 10.1534/g3.120.401491

Kessner D, *et al*. (2013). Maximum likelihood estimation of frequencies of known haplotypes from pooled sequence data. Molecular Biology and Evolution 30(5): 1145–1158. doi: 10.1093/molbev/mst016

Li H. (2013). Aligning sequence reads, clone sequences and assembly contigs with BWA-MEM. arXiv preprint, 1303-3997.

Li H. (2018). Minimap2: pairwise alignment for nucleotide sequences. Bioinformatics 34(18): 3094-3100. doi: 10.1093/bioinformatics/bty191

Long Q, *et al*. (2011). PoolHap: Inferring haplotype frequencies from pooled samples by next generation sequencing. PloS ONE 6(1): e15292. doi: 10.1371/journal.pone.0015292

Lu F, *et al*. (2013). Switchgrass genomic diversity, ploidy, and evolution: novel insights from a network-based SNP discovery protocol. PLoS Genetics 9(1): e1003215. doi: 10.1371/journal.pgen.1003215

Manching H, *et al*. (2017). Phased Genotyping-by-Sequencing enhances analysis of genetic diversity and reveals divergent copy number variants in maize. G3 Genes|Genomes|Genetics, 7(7): 2161-2170. doi: 10.1534/g3.117.042036

Nagy I, *et al.* (2022). Chromosome-scale assembly and annotation of the perennial ryegrass genome. Submitted for publication.

O'Neil ST and Emrich SJ. (2012). Haplotype and minimum-chimerism consensus determination using short sequence data. BMC Genomics 13(Suppl 2): S4. doi: 10.1186/1471-2164-13-S2-S4

Page JT, *et al*. (2014). BamBam: genome sequence analysis tools for biologists. BMC Research Notes 7: 829. doi: 10.1186/1756-0500-7-829

Patterson M, *et al*. (2015). WhatsHap: Weighted Haplotype Assembly for Future-Generation Sequencing Reads. Journal of Computational Biology. 22(6): 498-509. doi: 10.1089/cmb.2014.0157

Scheet P and Stephens M. (2006). A fast and flexible statistical model for large-scale population genotype data: Applications to inferring missing genotypes and haplotypic phase. The American Journal of Human Genetics 78(4): 629-644. doi: 10.1086/502802

The Potato Genome Sequencing Consortium. (2011). Genome sequence and analysis of the tuber crop potato. Nature 475**,** 189–195. doi:10.1038/nature10158

Tinker NA, *et al*. (2016). Haplotag: Software for Haplotype-Based Genotyping-by-Sequencing Analysis. G3 Genes|Genomes|Genetics 6(4): 857-863. doi: 10.1534/g3.115.024596

Untergasser A., *et al*., (2012). Primer3—new capabilities and interfaces. Nucleic Acids Research 40(15), e115. doi: 10.1093/nar/gks596

Verwimp C, *et al*., (2018). Temporal changes in genetic diversity and forage yield of perennial ryegrass in monoculture and in combination with red clover in swards. PLoS ONE 13(11): e0206571. doi: 10.1371/journal.pone.0206571

Wong T, *et al*. (2011). HaploJuice : accurate haplotype assembly from a pool of sequences with known relative concentrations. BMC Bioinformatics 19(1): 389. doi: 10.1186/s12859-018-2424-7

Zagordi O, *et al*. (2011). ShoRAH: estimating the genetic diversity of a mixed sample from next-generation sequencing data. BMC Bioinformatics 12(1): 119. doi: 10.1186/1471-2105-12-119

Zhang J, *et al.* (2014). PEAR: A fast and accurate Illumina Paired End reAd mergeR. Bioinformatics 30(5): 614–620. doi: 10.1093/bioinformatics/btt593
